# Modeling collagen fibril degradation as a function of matrix microarchitecture

**DOI:** 10.1101/2024.08.10.607470

**Authors:** B. Debnath, B. N. Narasimhan, S. I. Fraley, P. Rangamani

## Abstract

Collagenolytic degradation is a process fundamental to tissue remodeling. The microarchitecture of collagen fibril networks changes during development, aging, and disease. Such changes to microarchitecture are often accompanied by changes in matrix degradability. *In vitro*, collagen matrices of the same concentration but different microarchitectures also vary in degradation rate. How do different microarchitectures affect matrix degradation? To answer this question, we developed a computational model of collagen degradation. We first developed a lattice model that describes collagen degradation at the scale of a single fibril. We then extended this model to investigate the role of microarchitecture using Brownian dynamics simulation of enzymes in a multi-fibril three dimensional matrix to predict its degradability. Our simulations predict that the distribution of enzymes around the fibrils is non-uniform and depends on the microarchitecture of the matrix. This non-uniformity in enzyme distribution can lead to different extents of degradability for matrices of different microarchitectures. Our model predictions were tested using *in vitro* experiments with synthesized collagen gels of different microarchitectures. Experiments showed that indeed degradation of collagen depends on the matrix architecture and fibril thickness. In summary, our study shows that the microarchitecture of the collagen matrix is an important determinant of its degradability.

## 1 Introduction

Collagen is the most abundant protein present in tissues. Enzymatic degradation of collagen is an important process in both physiological and pathological conditions (Wohlgemuth et al., 2023; Gupta et al., 2024). For instance, an imbalance between collagen degradation and production can lead to increased collagen accumulation resulting in fibrosis (Wynn and Ramalingam, 2012; McKleroy et al., 2013). Such fibrotic environments have been associated with an impaired degradative environment (McKleroy et al., 2013). Studies have shown that the microarchitecture of collagen in the fibrotic extracellular matrix (ECM) is substantially different from the healthy tissues and they show higher resistance to degradation (Huang et al., 2009; Philp et al., 2018). As another example, cancer-associated fibroblasts in the tumor microenvironment secrete collagen and continuously remodel their matrix. This abnormal remodeling can lead to the cancerous microenvironment possessing a higher density of collagen and very different microarchitecture than a healthy ECM (Nebuloni et al., 2016). A body of work argues that alterations in collagen degradation can result in metastasis (Liotta et al., 1980; Winkler et al., 2020), and suggests possible connections between the matrix microarchitecture and its degradability (Dewavrin et al., 2014; Ranamukhaarachchi et al., 2019; Ashworth and Cox, 2024; Narasimhan and Fraley, 2024). However, the role of matrix microarchitecture in determining the degradability at the cellular length scale remains an open question. Here, we used multi-scale modeling and *in vitro* experiments to investigate possible mechanisms of matrix degradation at this length scale.

Most models of collagen gel degradation use Michaelis-Menten kinetics to model the collagenase activity and show good agreement with experiments in predicting the rate of total mass loss (Tzafriri et al., 2002; Metzmacher et al., 2007; Ray et al., 2013; Vuong et al., 2017). These models cannot address the connection between the microarchitecture and degradation because of their continuum structure. Another class of models have treated collagen fibril as a thin filament of negligible thickness (diameter ∼ 10−20 nm) and proposed a degradation mechanism of one filament based on movements of a few collagenase molecules over the filament surface (Saffarian et al., 2004; Sarkar et al., 2012). However, many experimental findings have reported that the collagen fibrils in a matrix are significantly thicker (diameter in the range ∼ 0.1 − 0.5 *µ*m) than the value used in these models (Bhole et al., 2009; Flynn et al., 2013; Staunton et al., 2016; Ranamukhaarachchi et al., 2019; Seo et al., 2020). Therefore, an open challenge in the field is to determine how a filament-scale model can be extended to a fibril-scale model, and subsequently, to a model where multiple fibrils are interacting with many collagenase molecules in a three-dimensional matrix.

In this work, we investigated how matrix microarchitecture can affect its degradability. We developed a fibril-scale model using lattice-based approach to predict the degradation of single fibril. To predict degradation of multiple fibrils by many enzyme molecules in a matrix, we used Brownian dynamics to capture the enzyme distribution surrounding the fibrils in a three-dimensional matrix environment. We then combined the fibril-scale model with the enzyme distribution obtained from the Brownian dynamics simulations to predict the degradation of collagen matrices. Our model predicted that differences in the microarchitecture between two matrices of same collagen density can lead to different extents of degradation. Additionally, we predicted that fibril thickness can be an important determinant of degradation. We tested this prediction against *in vitro* experiments using collagen gels of different architectures, which showed that indeed, fibril degradation depends on the matrix microarchitecture. Thus our study may provide new insights for understanding matrix alterations associated with disease and may influence the development of matrix targeted therapeutics, biomaterials and controlled drug delivery (Daly et al., 2020; Kim et al., 2022; Gupta et al., 2024).

## 2 Methods

In this section, we describe the details of the single fibril model, Brownian dynamics (BD) simulations, and experimental methods. We elaborate on each step of the model development and justify the assumptions based on previous experimental observations. The notation and symbols used in this work are shown in Table 1.

**Table 1:** Notation and list of symbols.

| Notation | Description |
| --- | --- |
| $d_f$ | mean diameter of a fibril |
| $\ell_f$ | mean length of a fibril |
| $d_{TC}$ | diameter of the tropocollagen unit |
| $\ell_{TC}$ | length of the tropocollagen unit |
| $d_E$ | mean hydrodynamic diameter of an enzyme |
| $d_m$ | length of one lattice site |
| $R$ | regular site |
| $V$ | vulnerable site |
| $EE$ | enzyme with two domains |
| $E_*E_*$ class of symbols | enzyme partially ($E_*E$) or, fully bound ($E_*E_*$)<br>to regular ($* = R$) or, vulnerable ($* = V$) sites |
| $k_*^*$ class of symbols | different rate constants |
| $N^s$ | total sites exposed on the fibril surface |
| $N_R^s$ | number of regular sites exposed on the fibril surface |
| $N_V^s$ | number of vulnerable sites exposed on the fibril surface |
| $N_{EE}$ | number of free enzymes |
| $N_{E_*E_*}^s$ class of symbols | number of enzymes partially (or, fully) bound<br>to regular (or, vulnerable) sites |
| $\phi$ | correction factor for intrinsic rates of full-length binding kinetics |
| $n_a$ | number of available lattice sites per enzymes at partially-bound state |
| $E_m$ | energy required to cross the symmetric energy barrier |
| $\lambda_m$ | extent of dissociation at the unwound state |
| $F$ | average force experienced by the lattice sites |
| $\gamma$ | surface energy per unit area |
| $\tau_c$ | characteristic time associated with force dependent kinetics |
| $\tau_e$ | time for disentanglement of tropocollagen chains |
| $l_p$ | nominal length scale |
| $N_e^0$ | number of enzymes surrounding a fibril |
| $(N_e^0)_{total}$ | total number of enzymes in the simulation box |
| $D_e$ | diffusivity of enzymes |
| $\phi_f$ | volume fraction of fibrils |
| $n_f$ | number of fibrils in the simulation box |
| $A_f^0$ | initial surface area of a fibril |
| $(A_f^0)_{total}$ | initial total surface area of all fibrils |

### 2.1 Model development for the degradation of single fibril

We consider a single collagen fibril of mean diameter *d*_*f*_ and mean length *𝓁*_*f*_. This fibril consists of a staggered arrangement of the triple helix tropocollagen units as shown in Fig. 1a (Orgel et al., 2006; Buehler, 2008; Marino and Vairo, 2014; Saini et al., 2020). The enzymatic degradation of the fibril occurs in the presence of collagenases such as matrix metalloproteinases (MMP). These collagenases cleave the triple helix at a site that is at a distance ∼ 67 nm (approx. 1*/*4 of the length of the triple helix) from the C-terminal of the tropocollagen unit (Fig. 1a) (Chung et al., 2004; Nagase and Visse, 2011).

**Figure 1:**
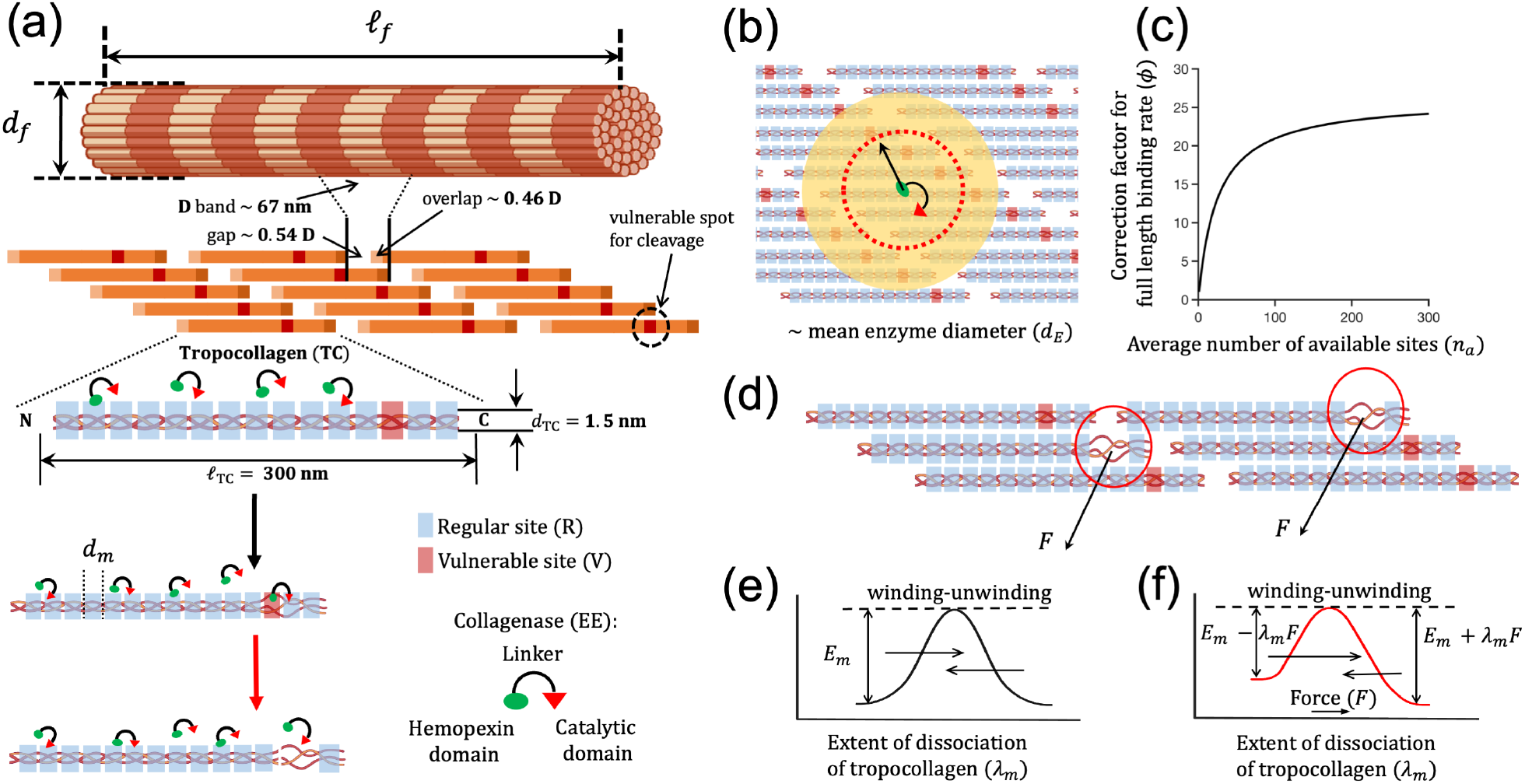
Modeling single fibril degradation. (a) Overview of the hierarchical organization of a collagen fibril and the degradation mechanism. We divide the tropocollagen unit into lattice sites (regular and vulnerable sites). The structure of the collagenase enzymes has two domains. (b) When one domain of the enzyme is bound to one lattice site, the other domain has access to only the neighboring unoccupied sites inside the red-dashed circle, whose radius is equivalent to the hydrodynamic mean diameter (*d*_*E*_) of the enzyme. (c) To incorporate this effect in the intrinsic rates of the full-length binding (when both domains of an enzyme are bound to the lattice sites)/hopping kinetics, we propose a correction factor (*ϕ*) as a function of available sites (*n*_*a*_). (d) We propose that the enzyme-induced irreversible unwinding of the vulnerable sites induces a force, which affects the unwinding rate of the neighboring regular sites. The symmetric (e) and asymmetric (f) energy barriers for the winding-unwinding process of a lattice site.

Previous experiments have reported that the collagenolytic degradation is a surface erosion process (Okada et al., 1992; Perumal et al., 2008; Tzafriri et al., 2002). The minimum effective pore size inside a collagen fibril is a few multiples of the diameter of the tropocollagen, which is *d*_TC_ ∼ 1.5 nm (Orgel et al., 2006). This dimension is smaller than the size of the collagenase molecules which is in the range 50-120 kDa with a mean hydrodynamic diameter of *d*_*E*_ ∼ 10 − 20 nm (Tyn and Gusek, 1990). Based on the dimensions and following previous studies (Okada et al., 1992; Perumal et al., 2008; Tzafriri et al., 2002), we assume that the diffusion of MMPs through the pores of the fibril can be ignored, and we model the fibril degradation as a surface erosion process.

To simulate the surface erosion of the fibril, we developed a lattice-based model. We divided the tropocollagen unit into lattice sites each of length *d*_*m*_. Based on the number of amino acid residues at the cleavage site region (Perumal et al., 2008), we set *d*_*m*_ = 8 nm. As the cleavage site is at a distance 1*/*4 of the length of the tropocollagen from its C-terminal, we assumed that there is one vulnerable site (V) per tropocollagen unit, and we treat the rest of the lattice sites as regular sites (R) (see Fig. 1a). With these dimensions, we developed a relation between the size of a fibril and the number of lattice sites available at its surface. To obtain the total number of lattice sites on the fibril surface, we next estimate the number of tropocollagen units on that surface. At any given time *t*, the number of tropocollagen units available for collagenolysis at the fibril surface is

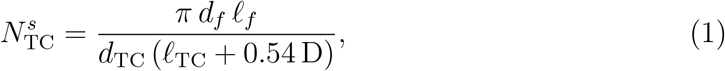

where *𝓁*_TC_ is the length of the tropocollagen unit and (0.54 D) is the gap between two tropocollagen units (D ∼ 67 nm) (see Fig. 1a). Thus, the total number of sites *N* ^*s*^ at time *t* exposed at the fibril surface is

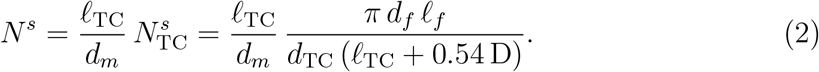

Eqn (2) highlights the relationship between the total number of sites available for degradation and the size of the fibril. The number of vulnerable sites 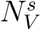 exposed at the surface is the same as 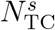 because one tropocollagen unit contains one vulnerable site, i.e.,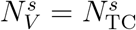. Therefore the remaining 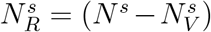 are regular sites. Using Eqn (2), we can estimate either *𝓁*_*f*_ or *d*_*f*_. As the vulnerable sites are not aligned at the same plane inside the fibril (see Fig. 1a) and the collagenolytic degradation is a surface erosion process, we treat *𝓁*_*f*_ as a constant and predict *d*_*f*_ from Eqn (2).

#### Development of a reaction scheme to model the loss of lattice sites on a fibril

Previous experimental findings have shown that the MMP class proteinases (for example, MMP1) hydrolyze the peptide bonds at the vulnerable site of the collagen type I (Chung et al., 2004; Nagase and Visse, 2011). Researchers hypothesized that the enzyme destabilizes the structure of the tropocollagen, and induces local unwinding around the vulnerable site before cleavage (Chung et al., 2004; Han et al., 2010; Perumal et al., 2008). Using the hypotheses of Chung et al. (2004) and Perumal et al. (2008), and making the following assumptions, we propose a reaction scheme for enzyme kinetics (see Fig. 1a and the supplementary information (SI) section SI1 (Fig. S1)):

1. There are two types of lattice sites on the collagen surface: regular site (R) and vulnerable site (V). The MMP has two domains: Hemopexin (Hpx) domain and Catalytic (Cat) domain connected by a linker (Visse and Nagase, 2003; Iyer et al., 2006). Manka et al. (2012) had shown a strict requirement of both domains to be linked together for efficient enzyme-collagen binding and collagenolysis.
2. Following Manka et al. (2012), we assume that the collagenase molecule has two domains, denoted by *EE*. Using these domains, the enzyme binds to the lattice sites via adsorption-desorption mechanism and hops on the sites. For the sake of simplicity, we assume that these two domains are equivalent and are of the same size.
3. First, using any one domain, the enzyme binds to a single site, in what is denoted as ‘reversible partial binding’. This type of binding can result in two possible reactions.

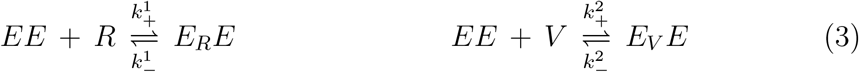

For the reversible partial binding (Eqn (3)), we implement the protein adsorptiondesorption kinetics (Adamczyk et al., 1999; Fang et al., 2005) (see a short note on adsorption kinetics in section SI2).
4. When one domain of the enzyme is bound to one site, the other domain can also bind to another site. When both domains of an enzyme are in bound state, we term this state as ‘full-length binding’. The enzyme molecule can jump or change track via hopping (Saffarian et al., 2004; Sarkar et al., 2012). It hops on the regular sites via reversible binding-unbinding of one domain to a regular site while its other domain remains bound to another regular site.

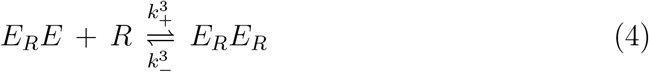
5. Once the enzyme reaches a vulnerable site with both domains in the bound state, hopping stops, and is followed by enzyme-induced unwinding and the formation of product sites. These product sites are no longer available for binding new enzyme molecules.

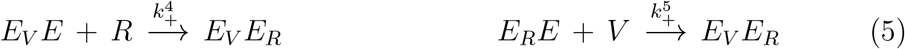

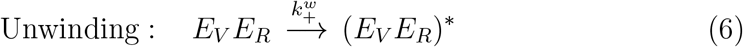

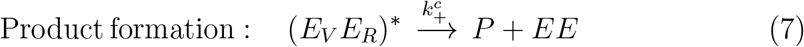

For the cases of full-length binding/hopping represented by Eqn (4)-(5), the forward reactions cannot be treated as second order kinetics because once one domain of an enzyme is bound to one site, its other domain does not have accessibility to all available lattice sites on the surface (see Fig. 1b). The unbound domain of the enzyme can find another lattice site for binding within a searching circle whose radius is equivalent to *d*_*E*_ (the mean hydrodynamic diameter of the enzyme). To correct the intrinsic rates of full-length binding kinetics, we multiply the rates with a correction factor *ϕ*. We propose a phenomenological function for *ϕ* as (Fig. 1c)

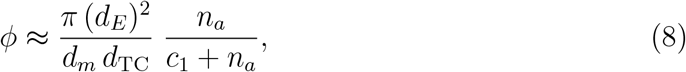

where *n*_*a*_ is the number of available sites on the fibril surface. See section SI3 for more details of *ϕ* and section SI4 for the system of ODEs representing Eqn (3)-(7).

The enzyme-induced permanent unwinding followed by cleavage happens only at the exposed vulnerable sites in collagenolysis. However, it is not clear how the fibril will lose other exposed regular sites at the surface due to the cleavage events so that the new surfaces can be easily accessible to the enzymes for further degradation (Chung et al., 2004; Saffarian et al., 2004; Nerenberg et al., 2008; Perumal et al., 2008; Sarkar et al., 2012). Hence, we propose that the enzyme-induced permanent unwinding at the vulnerable sites generates a force which assists the removal of more regular sites exposed at the surface. We use the nonequilibrium rate-process theory of Eyring (Eyring, 1935, 1936; Tobolsky and Eyring, 1943) and propose a new rate-term ℜ related to the force-dependent kinetics in an *ad hoc* manner in the system of ODEs. We briefly describe ℜ below.

As a result of thermal fluctuations, the lattice sites can be in any state between the triple helical and (temporary) unwound configurations (Perumal et al., 2008). If the rates of transition from the triple helical state to the unwound state and *vice-versa* are equal, there is no net change in the tropocollagen units in absence of enzymes. We hypothesize that enzyme-induced permanent unwinding that leads to cleavage generates a local stress (Fig. 1d). This mechanism is similar to the enzyme pulling chew-digest mechanism proposed by Eckhard et al. (2011). As a consequence, the other regular sites at the surface experience an external force which affects the kinetics by increasing their net rate of unwinding (Fig. 1e,f). This force can cause the slippage of chains, resulting in detachment from the surface (Adjari et al., 1994; Sung, 1995).

To incorporate the force-dependent kinetics, we add a new term 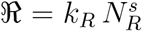 in an *ad hoc* manner to the system of ODEs (see section SI4 for the system of ODEs). Here *k*_*R*_ is the net rate of flow over the energy barrier towards force assisted unwinding (Fig. 1e,f). We propose the following expression for *k*_*R*_ (Glasstone et al., 1941; Stuart and Anderson, 1953)

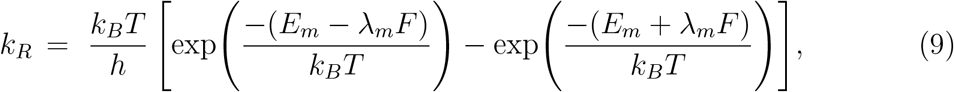

where *h* is Planck constant, *k*_*B*_ is the Boltzmann constant, and *T* is the temperature. Here, *E*_*m*_ is the energy required to cross the symmetric barrier (Fig. 1e), *λ*_*m*_ is the extent of dissociation in unwound state which is chosen as ∼ 3.6 Å (Perumal et al., 2008). Here *F* is the average force experienced by the remaining lattice sites. This force changes the energy barrier of unwinding to be asymmetric (Fig. 1f). If *F* = 0, *k*_*R*_ = 0. Following a heuristic approach, the average force (per remaining lattice site) is

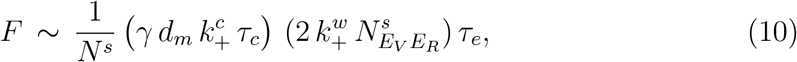

where 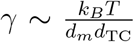 is the surface energy per unit area (Raphael and De Gennes, 1992; Léger and Creton, 2008), 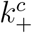 and 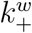 are rate constants of cleavage and enzymatic unwinding, respectively, 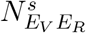 is the number of enzymes at fully-bound state (one regular and one vulnerable sites), *τ*_*c*_ and *τ*_*e*_ are, respectively, a characteristic time and time required to form disentangled chains during detachment from the surface. See sections SI5 for more details of *F, τ*_*c*_ and *τ*_*e*_, section SI6 for the reaction rate constants, and section SI7 for the initial conditions to solve the ODEs. We solved the system of ODEs using ODE23s in MATLAB (Mathworks, Natick, MA).

### 2.2 Brownian dynamics (BD) to estimate the enzyme distribution around fibrils in a matrix

We next set up a collagen matrix that consists of stationary non-overlapping cylindrical fibrils (Fig. 2a,b) for different fibril fractions. We modeled the enzymes as spheres with a diameter of 10 nm (Tyn and Gusek, 1990), and simulated their Brownian motion using overdamped Langevin dynamics (Ermak and McCammon, 1978). We used MATLAB to set up the simulations. The fibrils and enzymes are inserted in the simulation box such that they do not overlap and are treated as hard particles. The fibrils are randomly inserted using the methods given in Islam et al. (2016) and Ayad (2024). Periodic boundary conditions are applied in all three directions. For the sake of simplicity, all potential interactions among fibrils and molecules are neglected and the motion of the enzymes through the fibrous gel is simulated using Cichocki-Hinsen algorithm (Cichocki and Hinsen, 1990; Smith and Grima, 2017; Smith et al., 2017). The free diffusivity *D*_*e*_ of the enzyme is set to 74 *×* 10^−12^ m^2^/s (Schultz and Anseth, 2013). The time step for simulations is chosen Δ*t*_*s*_ = 10^−6^ s such that it is sufficiently small and the length increment in one time step is 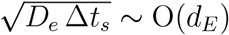, which is equivalent to the size of the enzyme *d*_*E*_ ∼ 10 nm. In simulations and model predictions, all length dimensions are scaled by *l*_*p*_ = 1 µm, and the dimensions of the simulation box are chosen as 5 *l*_*p*_ *×* 5 *l*_*p*_ *×* 5 *l*_*p*_. The entire codebase used in this work along with a readme file is available at https://github.com/RangamaniLabUCSD/Collagen_matrix_degradation.git.

**Figure 2:**
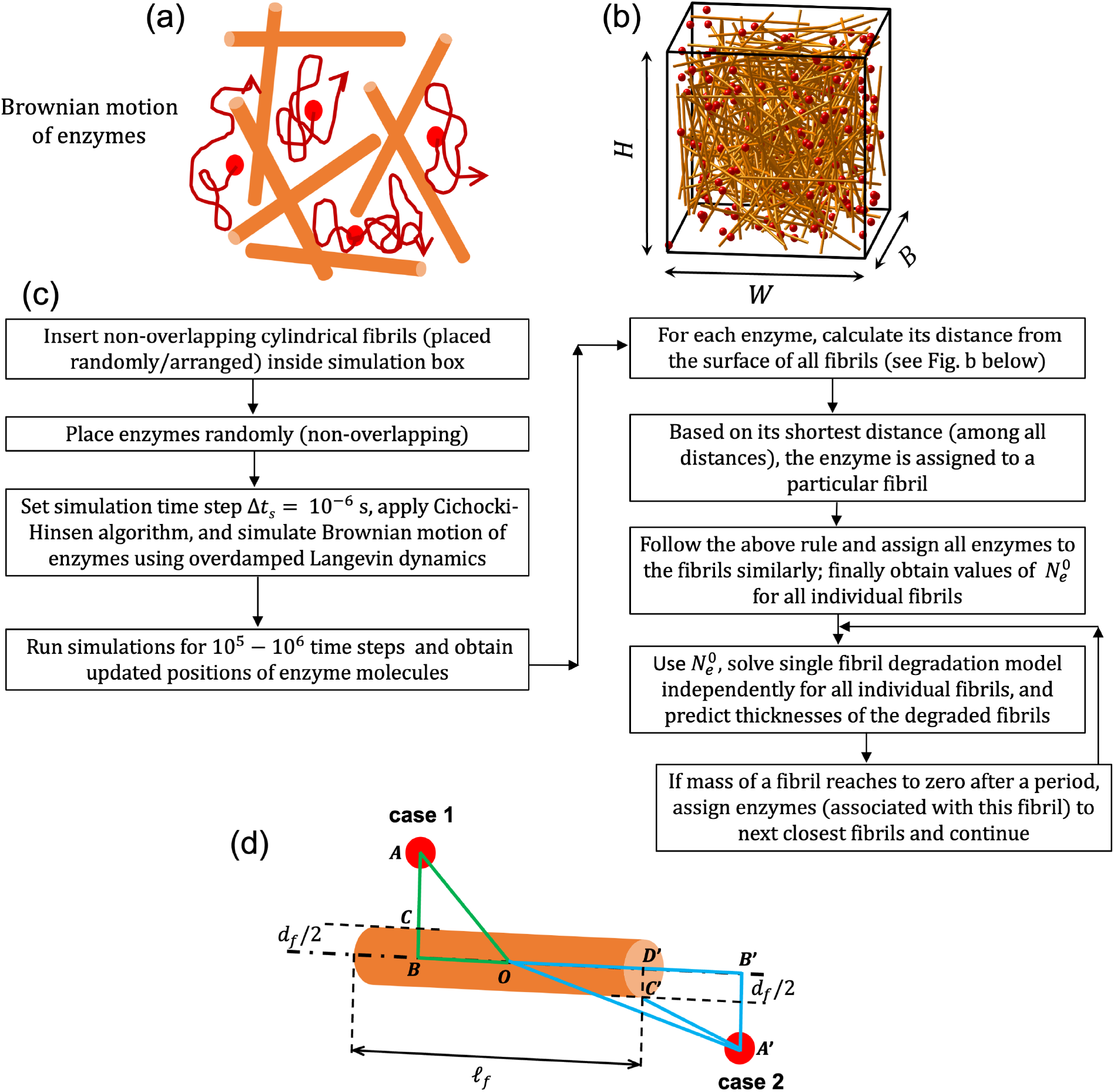
Brownian dynamics and hybrid simulation. (a) Schematic of the Brownian motion of enzymes inside the pores. (b) Configuration of the Brownian dynamics simulation where the cylinders represent collagen fibrils and the red spheres represent the enzyme molecules (not to scale). (c) Flowchart of the hybrid modeling approach. (d) Schematic representing the distance of an enzyme from the surface of a fibril. Based on the position of the enzyme with respect to any fibril surface, two cases are possible. For case 1, the distance chosen is *AC* = *AB* − *d*_*f*_ */*2. Otherwise, 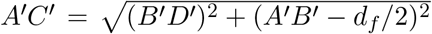 is the chosen distance for case 2, where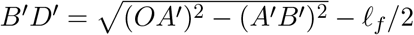.

For a length scale *l*_*p*_ = 1 µm and the diffusivity of the enzyme (collagenase) *D*_*e*_ ∼ 10^−10^ m^2^/s (Schultz and Anseth, 2013), the time scale is of 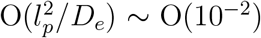. Hence the limit of instantaneous diffusion is valid in the long time limit for the chosen simulation box dimensions (Tzafriri et al., 2002). Based on this assumption, we proposed a hybrid framework (Fig. 2c). In this framework, we run the simulation for a period of time and then obtain the number of enzymes 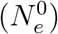 surrounding the fibrils based on the shortest distances of enzymes from the fibrils (Fig. 2d). We use the values of 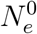 to solve the single fibril model where the fibril-scale model considers the binding-unbinding and other types of interactions among the enzymes and fibrils through a set of reactions. Using this hybrid modeling framework, we predict the degradation of all individual fibrils, effectively the degradation of a matrix.

To perform the BD simulations, we chose an enzyme concentration ∼ 2.5 µg/mL. For the simulation box volume 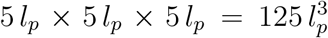, the weight of enzymes is 312.5 *×* 10^−18^ g. The molecular weight 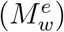 of collagenase varies between 70-130 kDa. We have chosen 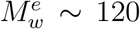 kDa (Eckhard et al., 2011). For this chosen value, the estimate of total number of enzymes in simulation volume is 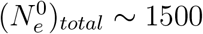.

We fixed the enzyme concentration and vary the collagen concentration, or equivalently, the fibril fraction *ϕ*_*f*_. Here *ϕ*_*f*_ is the ratio of the volume of all fibrils to the volume of the simulation box. See section SI8 for more details on estimation of the fibril fraction *ϕ*_*f*_ from a collagen concentration. By varying fibril diameter *d*_*f*_, fibril length *𝓁*_*f*_ and number of fibrils *n*_*f*_, we vary *ϕ*_*f*_ in the range 0.003 − 0.03, i.e. 0.3 − 3% for organs such as brain, liver, kidney, etc. (Tarnutzer et al., 2023). These three parameters, *d*_*f*_, *𝓁*_*f*_ and *n*_*f*_, are important factors to address different matrix microarchitectures. The orientation and curvature (or, crimping) of the fibrils can be other factors (Ashworth and Cox, 2024), however, we did not consider their roles in the present work.

### 2.3 Experimental methods

Following the experimental methodologies of Ranamukhaarachchi et al. (2019), we used rat tail type I collagen to synthesize the collagen gels. To vary the microarchitecture of the gels, we used polyethylene glycol (PEG) as a macro-molecular crowding agent. After polymerization of collagen using PEG at 37°C, PEG was washed out of the gels by rinsing them with Dulbecco’s Modified Eagle Medium (DMEM) solution. We then treated the gels with a bacterial collagenase. To characterize the microarchitectures of the gels preand post-degradation, we performed fast green staining and imaging using the confocal fluorescence microscopy and scanning electron microscopy. See section SI9 for more details of the experimental methods.

## 3 Results

### 3.1 Single fibril model captures experimentally observed degradation rates

We first validated the reaction scheme for a single fibril surface erosion model against previously published experiments of Flynn et al. (2013). In the absence of external loading, our model captured the experimental trend (Fig. 3a). See section SI10 for more details related to the validation procedure. When the fibril is under external loading, perhaps the external tension increases the stability of the triple helices by increasing the energy barrier for enzymatic unwinding (Chang et al., 2012; Chang and Buehler, 2014; Tonge et al., 2015; Saini et al., 2020; Topol et al., 2021), which is different from the mechanism for an isolated triple helix under external tension (Adhikari et al., 2011, 2012). Under low loading of ∼ 2 pN per tropocollagen monomer (Flynn et al., 2013; Tonge et al., 2015), a small increase in the energy barrier *E*_*m*_ for enzymatic unwinding by *δE*_*m*_ = 0.013 *E*_*m*_ explains the experimental trend satisfactorily (Fig. 3a). See section SI10 for the choice of *δE*_*m*_ = 0.013 *E*_*m*_. Overall, our model captured the experimental trends for a degrading fibril under different external loading conditions reasonably well.

**Figure 3:**
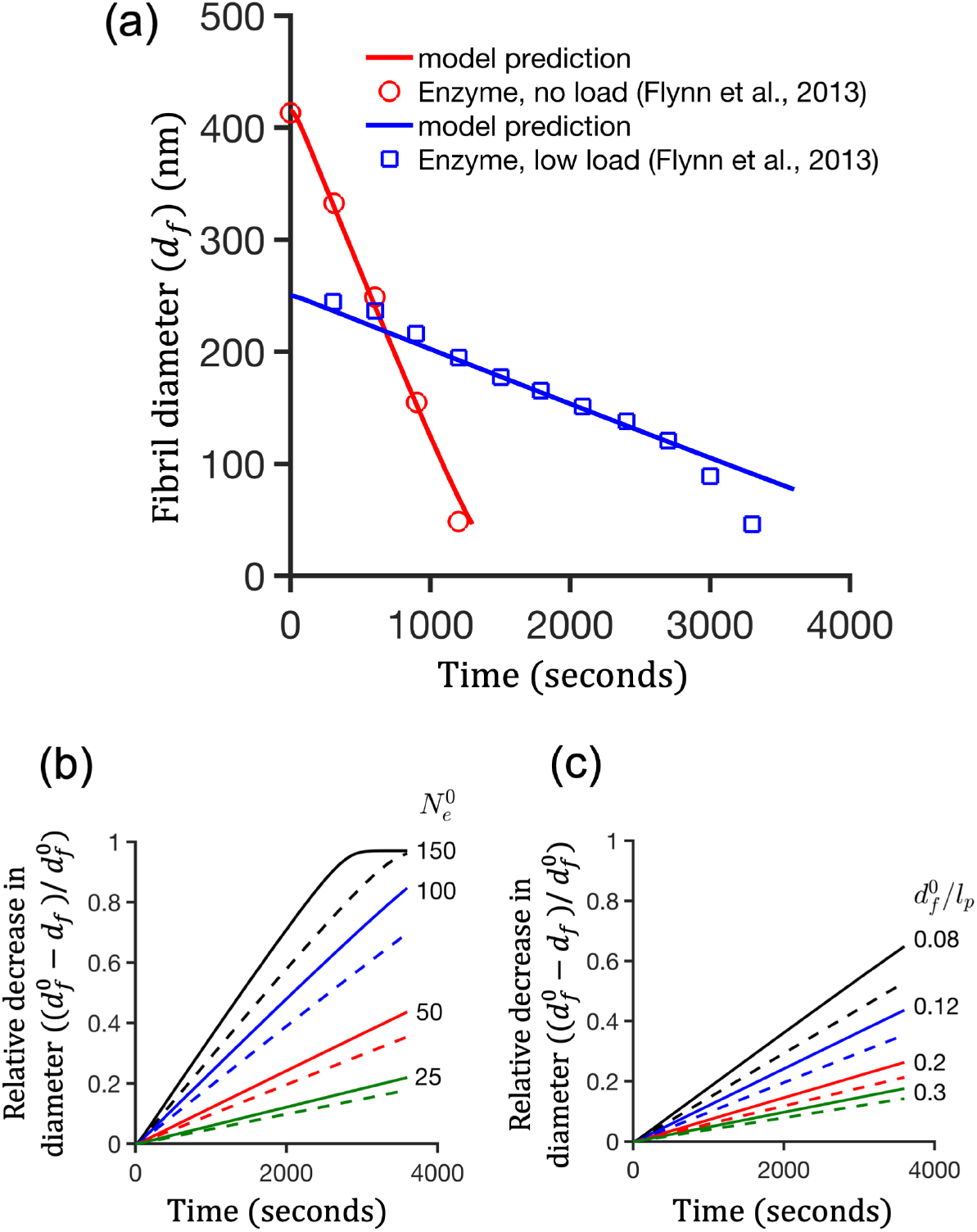
Degradation of single fibril. (a) Validation of the single fibril model with the experimental findings of Flynn et al. (2013). Relative decrease of fibril diameter 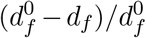 or, the extent of degradation) with time for two cases: (b) fixed initial fibril diameter 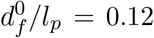 and varying number of enzymes surrounding a fibril 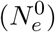; (c) Varying 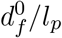 and fixed 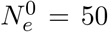. Here *l*_*p*_ = 1 µm is a nominal length scale. The length of the fibril is *𝓁*_*f*_ */l*_*p*_ = 2 in (b) and (c). The solid and dashed curves in (b) and (c) correspond to 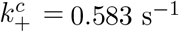 (Mallya et al., 1992) and 0.472 s^−1^ (Welgus et al., 1982), respectively.

In a surface erosion process, the degradability of a fibril must be proportional to the ratio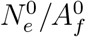, where 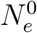 and 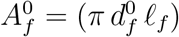 are the number of enzymes surrounding a fibril and the initial surface area of the fibril, respectively. For a fibril of fixed initial diameter 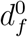, an increase in 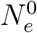 increases the degradability (Fig. 3b). However, ‘for a fixed value of 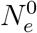, the degradability decreases with the increase in 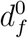 (Fig. 3c). The quantity 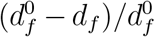 represents the relative decrease in the diameter. Its smaller value represents less degradation and *vice versa*. We provided a few results (Fig. S2) on degradability and the ratio 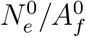 in section SI10. In summary, our model predictions captured the scaling related to a surface erosion process.

### 3.2 Degradation of collagen matrices

Having established that the single fibril model captures experimentally observed degradation rates (Fig. 3a), we next focused on degradation of collagen fibrils in matrices. Before addressing matrix degradation, we define two important dependent parameters: the initial volume fraction of fibrils as 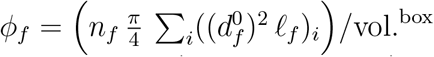, and the initial total surface area (scaled) of fibrils as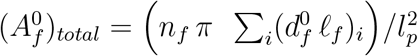.

#### Degradation of matrices with uniform fibrils

We first considered uniform fibrils of same initial diameter 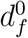 and length *𝓁*_*f*_, and investigated the effect of the number of fibrils 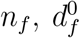, and *𝓁*_*f*_ on degradation. Using the hybrid approach described in Fig. 2b,c, we counted the number of enzymes 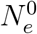 surrounding every single fibril (Fig. 4a). We used this count of the number of enzymes to obtain a probability density function (PDF) of 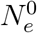 for a given matrix (Fig. 4b).

**Figure 4:**
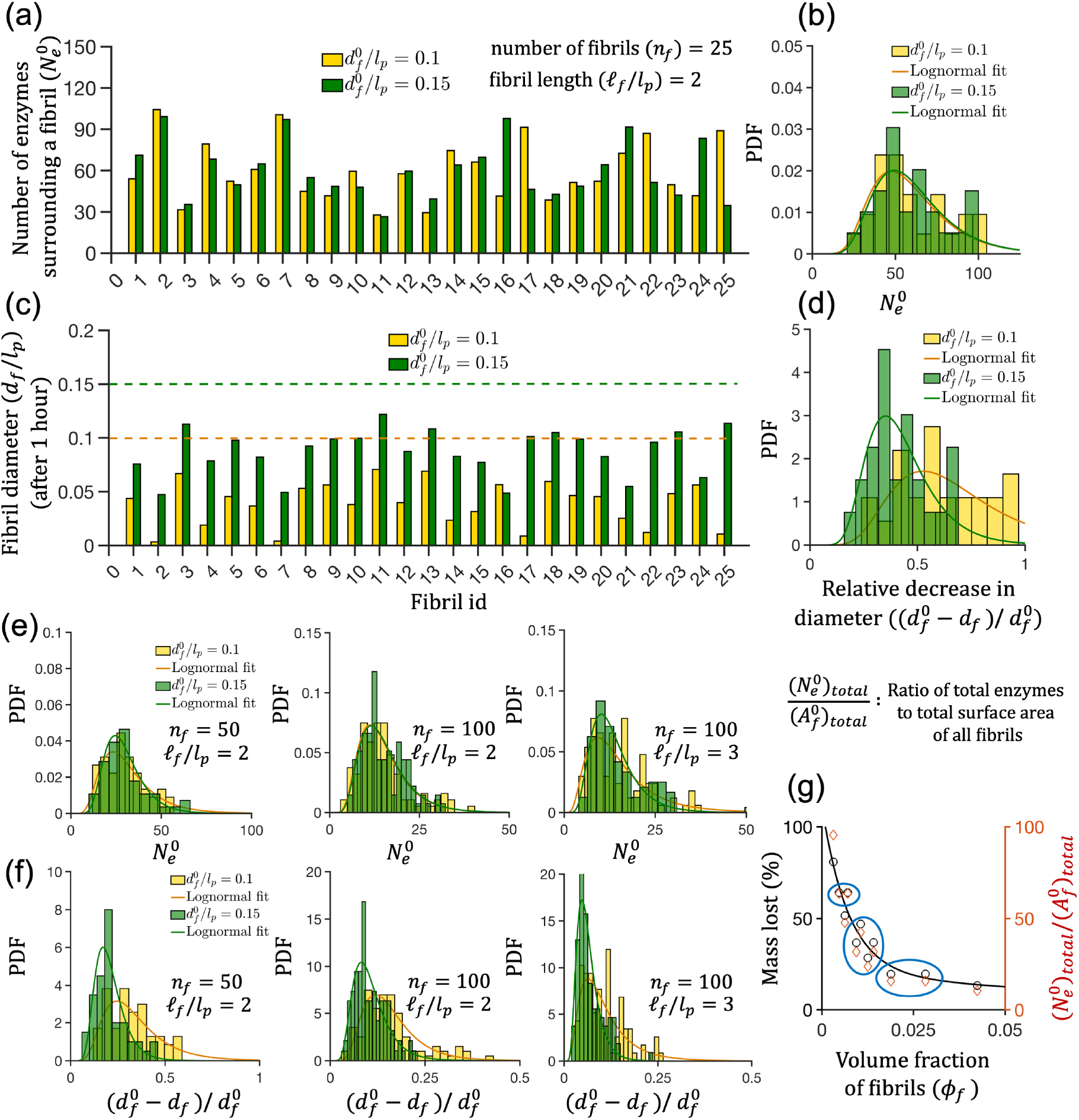
Degradation of collagen matrices. (a) The values of the number of enzymes 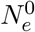 surrounding the individual fibrils in a matrix for two different matrices of same fibril length *𝓁*_*f*_ and number of fibrils *n*_*f*_, but for two different values of initial fibril diameter 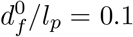 and 0.15. (b) The probability density function (PDF) of 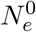 generated using the data in (a). (c) The predicted diameters of the fibrils after 1 hour of degradation by enzymes for the cases in (a). (d) The PDFs representing the relative decrease in diameter 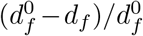 for the data shown in (c). Panels (e) and (f) represent PDFs of 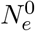 and 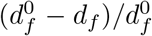 for different values of 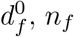 and *𝓁*_*f*_. (g) The percentage of the mass lost (left ordinate) and the ratio 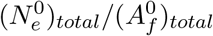 (right ordinate) with the volume fraction of fibrils *ϕ*_*f*_. The black curve in (g) represents a fit to the data points corresponding to the left ordinate. The data points inside the blue circles show non-monotonous trends of the mass lost and the ratio 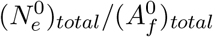 with respect to *ϕ*_*f*_.

We observed that the enzyme distribution was not uniform across the fibrils in a matrix (Fig. 4a,b), suggesting that the organization of the collagen fibrils was a major determinant of the enzyme distribution. There are a few factors that can result in this non-uniform distribution of enzymes around fibrils. First, the cylindrical geometry of fibrils can result in an anisotropy of the diffusive motion of enzymes (Stylianopoulos et al., 2010; Chen et al., 2021). Even if the cylindrical geometries of the fibrils are taken into account, for enzyme distributions in a matrix to be nearly uniform, fibrils should be organized in an equidistant manner with parallel alignment. This organization ensures a uniform pore size around cylindrical fibrils and can result in nearly uniform enzyme distribution (Fig. S3). However, the random placement and random orientation of fibrils induces a non-uniform distribution of pore sizes. These factors together can result in a non-uniform enzyme distribution in a fibrillar matrix. Our simulations show that this finding of non-uniform distribution of enzymes holds for different values of fibril diameter, fibril length, and number of fibrils (Fig. 4e).

The enzyme distributions do not differ much between two matrices if we vary only 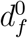, fixing *𝓁*_*f*_ */l*_*p*_ = 2 and *n*_*f*_ = 25 (Fig. 4b). Using the values of the number of enzymes in Fig. 4a, we solved the single fibril model for each of the individual fibrils and calculated the diameters of the fibrils (Fig. 4c). As expected, each fibril degrades to a different extent because of the difference in the number of enzymes surrounding it. Using these single fibril data, we obtained the PDF of the relative decrease in the diameter of the fibril 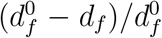(Fig. 4d). Thus, we find that although the initial fibril diameter is uniform, the degraded fibrils have a distribution of diameters because of the nonuniform enzyme distribution.

We next investigated the effect of the number of fibrils *n*_*f*_ on the extent of degradation. From our simulations, we find that the enzyme distribution is a strong function of *n*_*f*_ (Fig. 4e). For a fixed value of 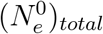, we can expect a decrease in the average value of 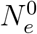 per fibril if *n*_*f*_ increases. When comparing two matrices of the same 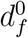 and *𝓁*_*f*_, the matrix with higher number of fibrils degrades to a lesser extent than the matrix with fewer fibrils (Fig. 4d,f). We also find that an increase in 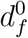 and *𝓁*_*f*_ decreases the degradability of the matrix for fixed values of *n*_*f*_ and 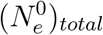(Fig. 4d,f) because o the decrease in the ratio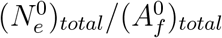.

#### Effect of matrix architecture on fibril degradation

Since we model the single fibril degradation as a surface erosion process, the extent of degradation for a matrix must be proportional to the ratio of the total number of enzymes to the total surface area of the fibrils 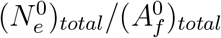. The overall mass lost, defined as

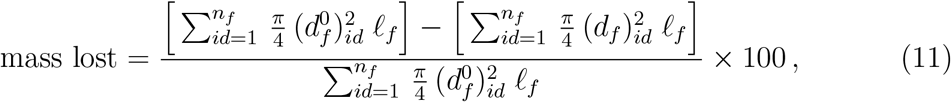

represents a direct estimate of the matrix degradation. For all combinations of 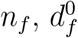 and *𝓁*_*f*_, we find a direct correlation between the mass lost and the ratio 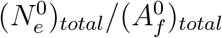 (Fig. 4g) when they are plotted against the fibril fraction *ϕ*_*f*_. However, the variation of the mass lost and 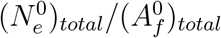 with *ϕ*_*f*_ are not strictly monotonous (blue circles in Fig. 4g), although their trends appear to be monotonically decreasing as *ϕ*_*f*_ increases. The circled data points in Fig. 4g show that for the same enzyme concentration, different extent of degradation can occur between two matrices of same fibril fraction *ϕ*_*f*_ (equivalently, the same collagen concentration) and *vice versa*. This finding leads to the following question: does the difference in microarchitecture of two matrices of same *ϕ* impact the extent of degradation? We answer this question by conducting the following simulations.

We fix the value of *ϕ*_*f*_ and the enzyme concentration, and consider the matrices with uniform fibrils. Without varying *𝓁*_*f*_, different matrices of same *ϕ*_*f*_ can be generated by adjusting the values of *n*_*f*_ and 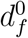 (Fig. 5a). For example, as 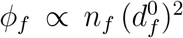, a 50% decrease in 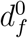 can increase *n*_*f*_ up to 4 times, resulting in twice the increase of the surface area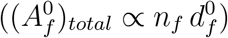. For the case of constant *ϕ*_*f*_ and smaller 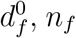 increases and shifts the enzyme distribution towards the left (Fig. 5b). Simulations show that a matrix composed of thinner fibrils degrades less than that a matrix composed of thicker fibrils (Fig. 5c). This is an unexpected result and counterintuitive to what we might expect from a single fibril model. In the single fibril degradation model, a thinner fibril degrades more than a thicker fibril if the enzyme concentration is the same (Fig. 3c) because of lesser surface area of a thinner fibril available to larger number of enzymes. In contrast, in a three-dimensional, randomly oriented and randomly placed fibril network, multiple factors affect the enzyme distribution as discussed previously. Thus, this finding highlights the importance of incorporating the three-dimensional spatial considerations including diffusion of enzymes and matrix microarchitectures into the model. The results in Fig. 5d-f show how the number of fibrils changes for different sets of 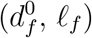 when *ϕ*_*f*_ is fixed, which directly influences the ratio 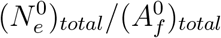, and the overall mass lost. In summary, our simulations predict that for the same enzyme concentration, uniform fibril diameter, and the same fibril fraction *ϕ*_*f*_, a matrix with thicker fibrils can degrade more than that with thinner fibrils (Figs. 5d-f).

**Figure 5:**
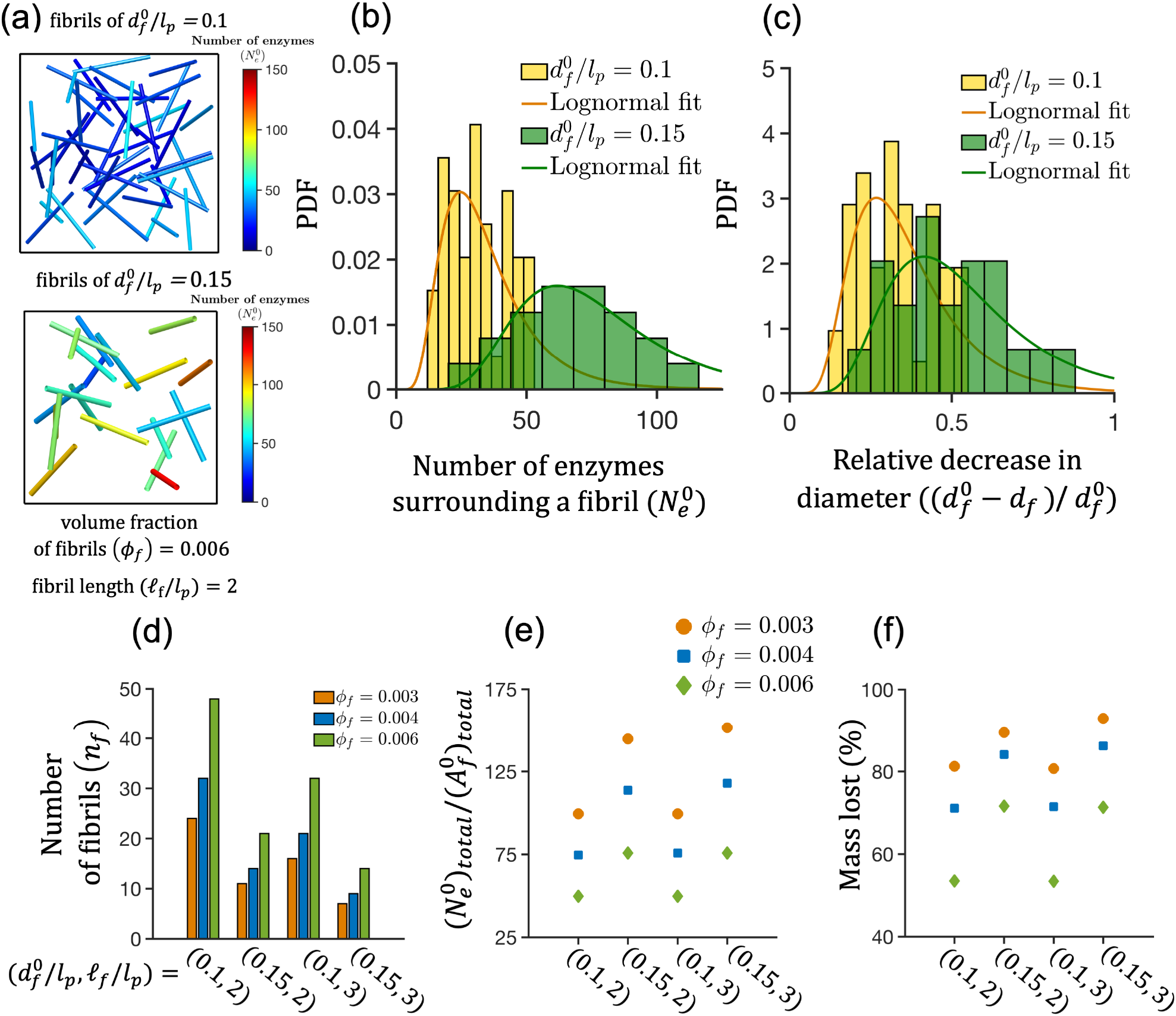
Matrix with thicker fibrils degrades more than that with thinner fibrils. (a) The initial configurations of two matrices having the same fibril fraction *ϕ*_*f*_ = 0.006. The color bars in (a) represent the number of enzymes 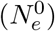 surrounding the fibrils. For the configuration in (a), the PDF of enzyme distribution 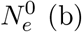 and the PDF of the extent of degradation of fibrils 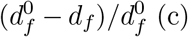. For matrices having different *ϕ*_*f*_, the first (d), second (e) and third (f) panels report the number of fibrils in matrices, 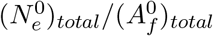 and the percentage mass lost (overall degradation of matrices), respectively. All results correspond to the total number of enzymes 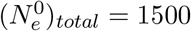 and time 60 minutes.

To compare this model prediction on matrix degradability, we performed *in vitro* experiments with synthetic collagen gels and investigated their degradability.

### 3.3 Experiments reveal that matrix microarchitecture governs degradability

Ranamukhaarachchi et al. (2019) previously showed that the microarchitecture of a collagen gel can be modulated using polyethylene glycol (PEG) as a macro-molecular crowding (MMC) agent. They also reported that the degradability of the gels varies with the matrix microarchitecture. Using this background, to test our model predictions, we prepared collagen gels with two different microarchitectures from the same collagen concentration. We used 2.5 mg/ml collagen (final concentration) and two different MMC concentrations: 2 mg/mL PEG designated as P2 and 8 mg/mL PEG denoted as P8 (Fig. 6a). We used a bacterial collagenase concentration of 2.5 µg/mL (final concentration) to perform degradation experiments. Using fast green staining images and scanning electron microscopy (SEM), we quantified the fibril length and thickness distributions respectively before and after degradation. We denote the gels post-degradation as P2x and P8x.

**Figure 6:**
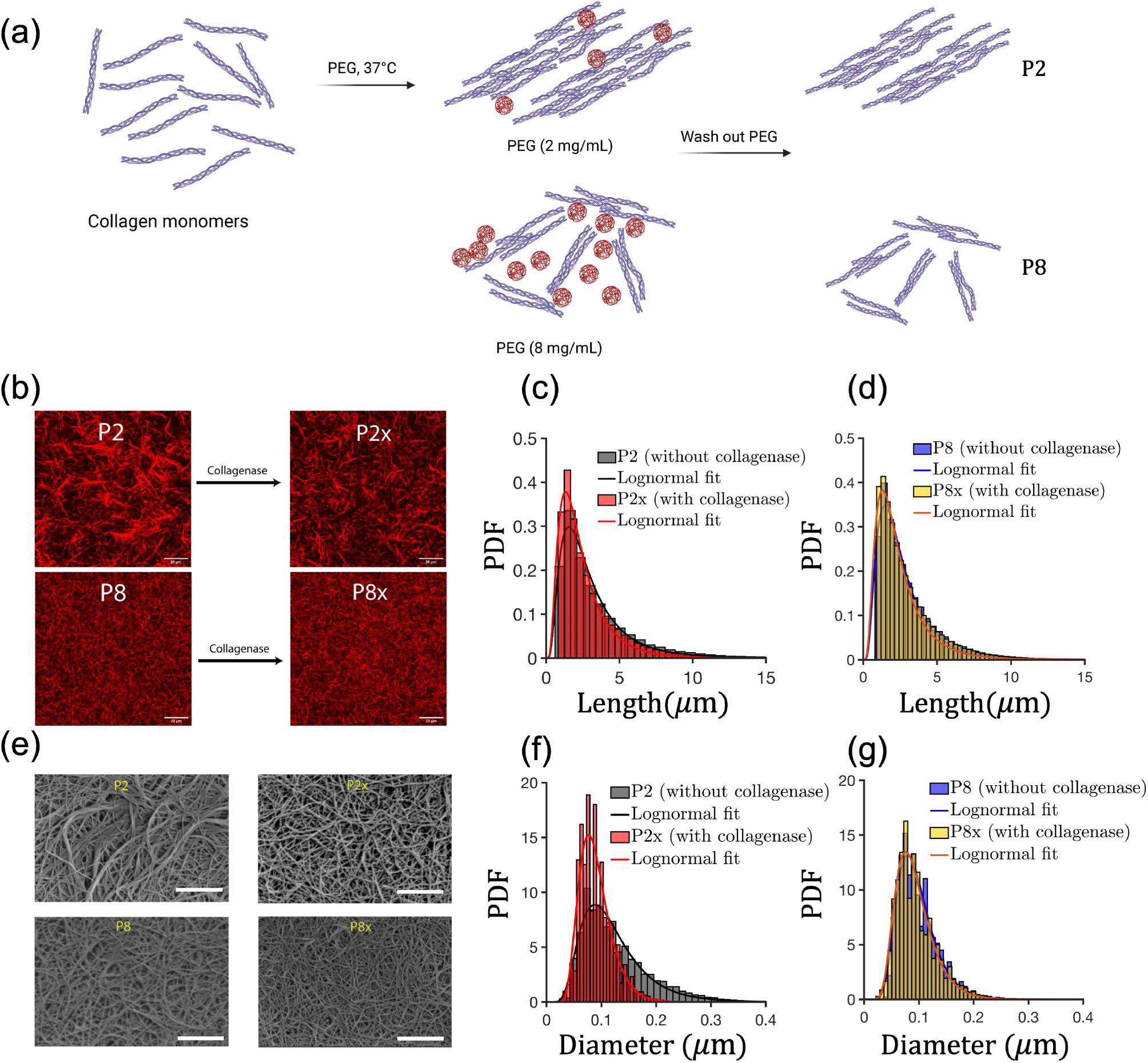
Experimental findings. (a) Preparation of collagen gels. (b) The fast green stained images of the gels (scale bar 20 *µ*m). The length distributions of (c) P2/P2x and (d) P8/P8x before and after treatment with collagenase from the green stained images. (e) The SEM images of P2/P2x and P8/P8x (scale bar 2 *µ*m). The histograms of fibril diameters for (f) P2/P2x and (g) P8/P8x.

Fig. 6b shows the fast green stained images, and the length distributions for P2/P2x (Fig. 6c) and P8/P8x (Fig. 6d). The mean length of the fibrils in P8 (∼ 3 *±* 0.2 µm) is slightly smaller than that of P2 (∼ 3.3 *±* 0.4 µm). From Fig. 6b, it is obvious that P2 degrades more than P8 as larger pores are present in P2x. However, there is no significant change in the length distributions before and after degradation in both P2/P2x and P8/P8x (Fig. 6c,d); the decrease in the mean length post-degradation is less than ∼ 8%. Hence the assumption of treating the fibril length *𝓁*_*f*_ as a constant in our model is supported by the experimental findings. The green stained images (Fig. 6b) reveal that the number of fibrils is higher in P8 than that in P2 (through visualization).

The SEM images (Fig. 6e) and the histograms of the fibril diameters (Fig. 6f,g) show that the fibrils are thicker in P2 than in P8. The mean diameter of the fibrils in P8 (∼ 0.09 *±* 0.02 µm) is ∼ 25-30% smaller than that of P2 (∼ 0.13 *±* 0.03 µm). After degradation, the decrease in the diameter occurs in both P2x and P8x (Fig. 6f,g). Quantification of the diameters of the fibrils show that P2 (Fig. 6f) degrades more than P8 (Fig. 6g). The decrease in the mean diameter is ∼ 30 − 40% in P2x and ∼ 15% in P8x. Thus, experiments validate our model predictions that a matrix with thicker fibrils can degrade more than one with thinner fibrils for the same collagen and collagenase concentrations.

#### Model predictions for matrices with non-uniform fibrils highlight the role of microarchitecture in degradation

To further reinforce the role of matrix microarchitecture, we note that the significant differences between the microarchitectures of P2 and P8 are primarily due to the number of fibrils and fibril diameter. In P2, there are less number of fibrils and the fibrils are thicker. However, in P8, there are higher number of fibrils but the fibrils are thinner, compared to P2 (Fig. 6). Our model predicts higher degradability of a matrix with thicker fibrils than that with thinner fibrils (Figs. 5). Thus, we can qualitatively explain why the matrix with the P2 architecture degrades more than the P8 architecture. However, our simulations in Fig. 5 are for fibrils of uniform initial diameter. Because the experimentally synthesized matrices had fibrils with a distribution of diameters, we next simulated the degradation of fibrils in matrices of experimentally-inspired fibril diameter distributions.

We generated the fibrils whose diameters matched the experimentally observed diameters of P2 and P8 (Fig. S4 and Fig. S5) while maintaining the fibril length to be the same (see section SI11 for more details). We used a fibril fraction of *ϕ*_*f*_ = 0.007 in both cases as this value is close to the collagen concentration used in the experiments. As a result, in these newly generated matrices, the number of fibrils is higher for P8 (*n*_*f*_ ∼ 75) than that in P2 (*n*_*f*_ ∼ 40) for the same fibril fraction *ϕ*_*f*_. We compared the outcomes of degradation in these conditions as shown in Fig. 7. The microarchitectures are different for P2 and P8 (Fig. 7a,b) in terms of the number of fibrils and the diameters. As a result, the enzyme distribution for P8 shifts to the left due to larger number of fibrils (Fig. 7c) and implies less number of enzymes per fibril. The diameter distributions (in the range 0.03-0.2 µm) of P2 and P8 before and after degradation (Fig. 7d,e) and the overall mass lost (Fig. 7f) indicate that P2 degrades more than P8 for the same collagen and enzyme concentrations, in agreement with experiments (Fig. 6e,f). Our simulation results using non-uniform fibril diameter only reinforce our model predictions that the matrices of the same collagen concentration can have very different microarchitectures and the matrices can degrade differently under the same collagenase concentration. In summary, our study reveals that matrix microarchitecture plays an important role in degradation of collagen matrices.

**Figure 7:**
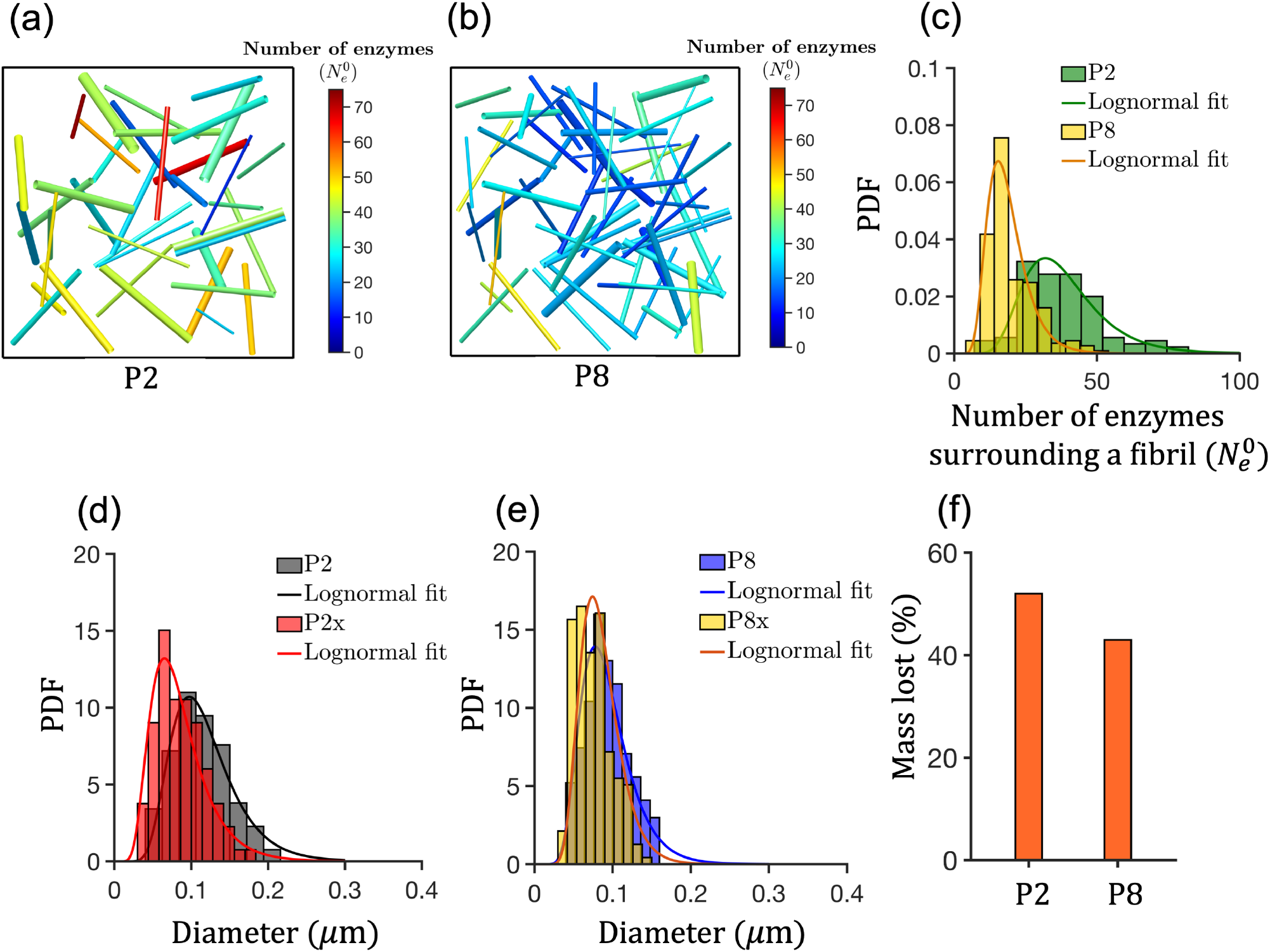
Simulations of matrices with non-uniform fibrils predict that P2 degrades higher than P8. The initial configurations of two matrices, P2 (a) and P8 (b), with the same fibril fraction *ϕ*_*f*_ = 0.007. The color bars in (a) and (b) represent the number of enzymes 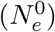 surrounding the fibrils. For the configuration in (a) and (b), the PDF of the enzyme distributions 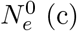, the PDF of the diameters of the fibrils for P2/P2x (d) and P8/P8x (e), and the overall mass lost (f). The results correspond to the total number of enzymes 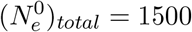 and time 60 minutes.

## 4 Discussion

In this study, using a combination of modeling and experiments, we investigated the collagenolytic degradation of collagen matrices. We showed that the matrix microarchitecture is a strong determinant of matrix degradability. The enzyme distribution in a matrix is not uniform due to the fibrillar network, and this enzyme distribution leads to fibrils of distributed diameters even if the fibrils are initially uniform in size. The matrices of the same collagen concentration can degrade differently under the same collagenase concentration because of the differences in the microarchitecture. These findings are backed by *in vitro* experiments with collagen gels of different microarchitectures.

Unraveling the connection between the matrix microarchitecture and degradation is important to design biomaterials and understand ECM remodeling at cellular length scale. The previous models are either continuum or discrete, and thus, they predict either overall rate of degradation or degradation at nanoscale (Tzafriri et al., 2002; Metzmacher et al., 2007; Vuong et al., 2017; Sarkar et al., 2012). The model and framework we developed in this work to investigate the collagen matrix microarchitecture and matrix degradation is novel in the sense that the previous models are not capable to address the connection between the degradation and microarchitectures. However, our model has also some drawbacks. The lattice-based single fibril model has a limitation to predict the spatial variations in the size and shape of a degrading fibril. In our simulations, the fibrils are non-stretchable and we did not consider explicitly the potential energy based attractive interactions among the fibrils and enzymes. At present, it is difficult to state *a priori* whether the potential interactions can affect the temporal evolution of the enzyme distribution which merits further attention.

In summary, our single fibril model and hybrid modeling framework effectively capture multi-scale effects to predict the degradation of three-dimensional matrices with different microarchitectures. Although relatively simple, this framework sheds light on how collagen matrix degradability is tuned by matrix microarchitecture. This has important implications for a number of fields, including matrix biology and biomaterials.

## Acknowledgements

This work was supported by a National Science Foundation grant DMS-1953469, American Cancer Society Research Scholar Grant RSG-21-033-01-CSM to S.I.F., the National Cancer Institute U54CA274502, a Prebys Research Heroes Grant to S.I.F. P.R. and S.I.F were also supported by the Wu Tsai Human Performance Alliance and the Joe and Clara Tsai Foundation. We would like to thank the UC San Diego School of Medicine Microscopy Core, which is supported by the National Institute of Neurological Disorders and Stroke grant P30NS047101.

## Supplementary Information (SI)

### SI1 Reaction scheme

**Figure S1:**
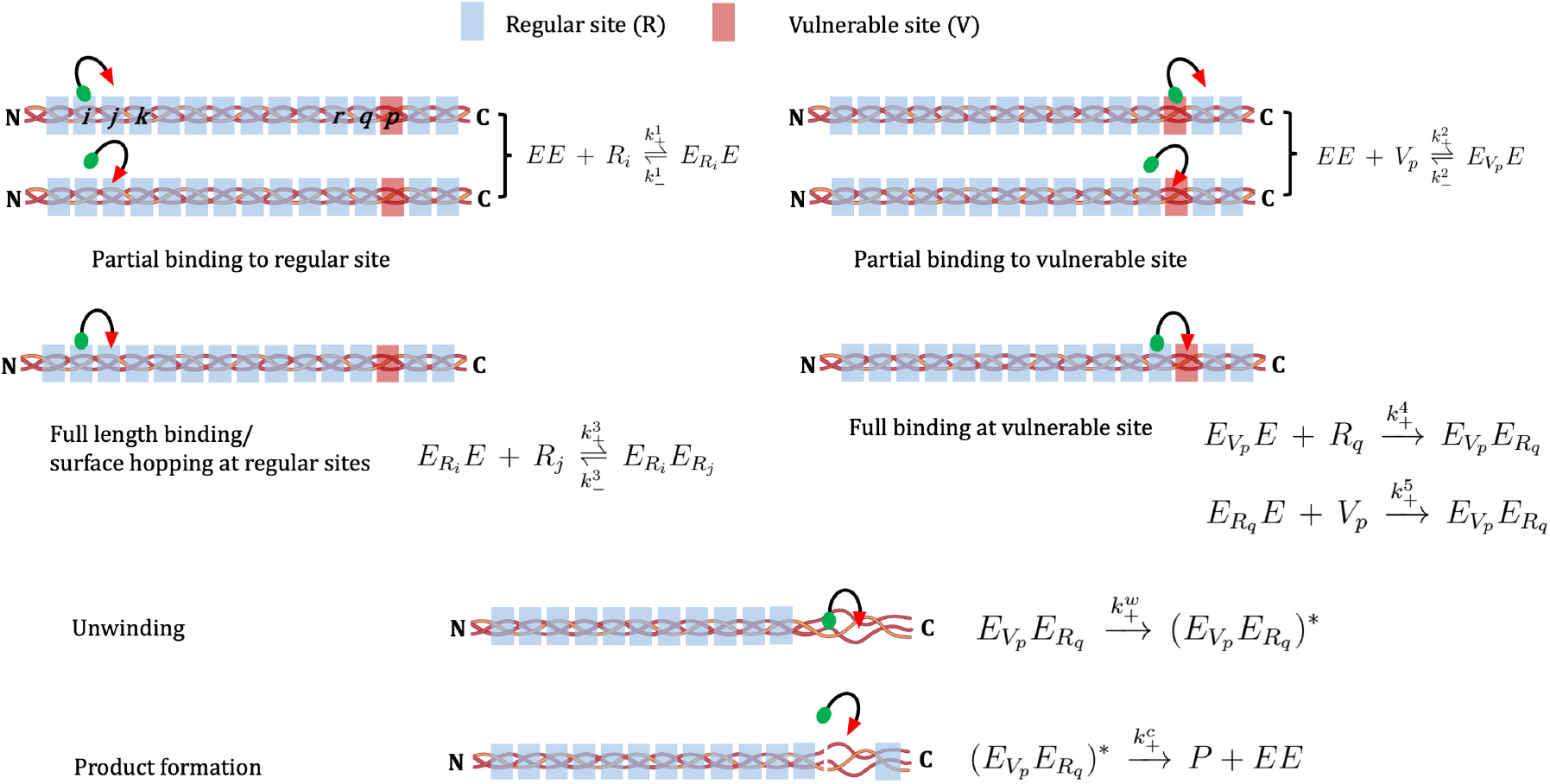
Schematic of the enzyme kinetics.

### SI2 Adsorption kinetics and Φ_*ad*_

For reversible partial binding (represented by Eqn (3)), we implemented the protein adsorption-desorption kinetics (Adamczyk et al., 1999; Fang et al., 2005). The rate of adsorption is proportional to the available surface function Φ_*ad*_ = 1 − (*θ/θ*_*max*_). This available surface function is related to the steric hindrance during adsorption (Adamczyk et al., 1999; Adamczyk, 2000; Fang et al., 2005), where *θ* and *θ*_*max*_ are the surface coverage fraction and its maximum saturated value, respectively. If the number of total enzymes is very small compared to the number of available sites, Φ_*ad*_(*θ*) → 1 in the limit of very low surface coverage. If the limit of low surface coverage does not hold true, some descriptions related to Φ_*ad*_(*θ*) are provided in earlier work (Schaaf and Talbot, 1989; Talbot et al., 2000; Adamczyk, 2000). In our work, the limit *N* ^*s*^ ≫ *N*_*EE*_(*t* = 0) holds true for low concentration of enzyme, where *N* ^*s*^ and *N*_*EE*_ are total number of sites exposed at the surface and number of enzymes, respectively. Thus we set Φ_*ad*_ = 1.

### SI3 Correction factor *ϕ* in the intrinsic rates of full-length binding kinetics

In our model, we treated the forward rates of Eqn (4)-(5) as pseudo first-order kinetics multiplied with a correction factor *ϕ*. The functional form of *ϕ* increases with the number (*n*_*a*_) of available lattice sites per enzyme at partially bound state. The value of *ϕ* is zero if there is no site available, *ϕ* = 1 when only one lattice site is available (*n*_*a*_ = 1), and *ϕ* must saturate around (*π* (*d*_*E*_)^2^)*/*(*d*_*m*_ *d*_TC_) which is equivalent to the number of lattice sites inside the searching radius *d*_*E*_ (see Fig. 1c). We proposed a phenomenological function for *ϕ* as

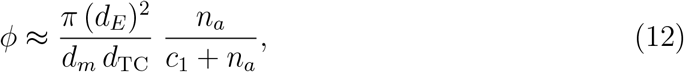

where 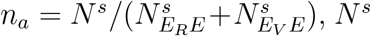 is the number of available sites on the fibril surface, 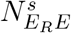 and 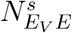 are the number of enzymes partially bound to one regular site and one vulnerable site, respectively. Here *c*_1_ = 25 is a dimensionless constant which is obtained by setting *ϕ*(*n*_*a*_ = 1) = 1 for *d*_*E*_ = 10 nm, *d*_TC_ = 1.5 nm and *d*_*m*_ = 8 nm.

### SI4 System of ODEs

The system of ODEs representing the reaction scheme Eqn (3)-(7) is the following

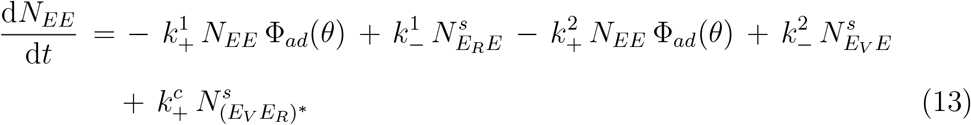

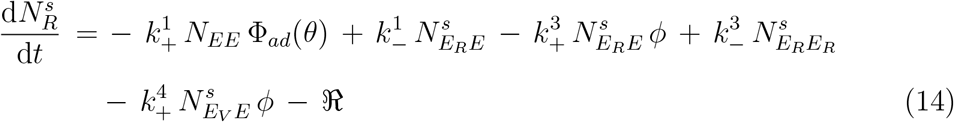

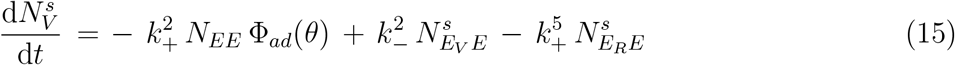

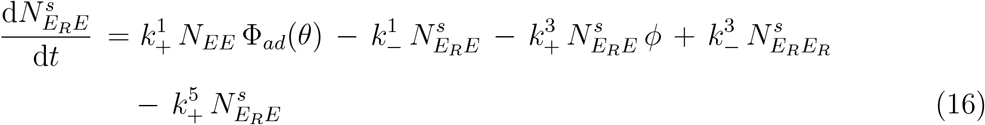

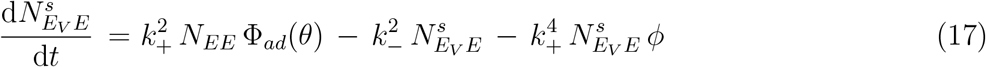

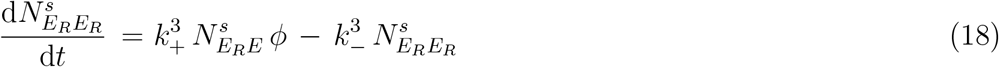

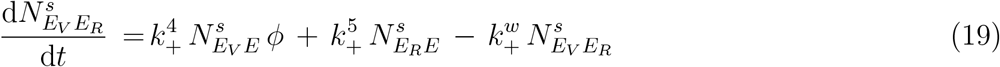

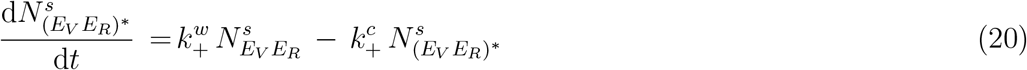

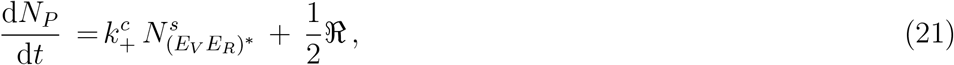

where 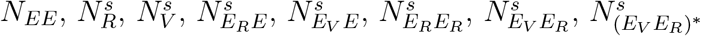 are numbers of free enzymes, regular sites, vulnerable sites, enzymes partially bound to one regular site, enzymes partially bound to one vulnerable site, enzymes fully bound to two regular sites, enzymes fully bound to one regular and one vulnerable site, enzymes fully bound to one regular and one vulnerable site (unwound state), and product sites, respectively. Here 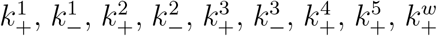 and 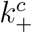 are the reaction rate constants having dimension [time]^−1^; Φ_*ad*_ and *ϕ* are available surface function and correction factor related to adsorption kinetics and full-length binding/hopping kinetics, respectively.

We included another term ℜ in the series of ODEs (Eqn (13)-(21)) in an *ad hoc* manner, which is related to force-assisted removal of regular sites. We discussed about ℜ in the main text and in the next section. The balances for the total number of enzymes and total number of sites are the following.

#### Balance for total number of enzymes

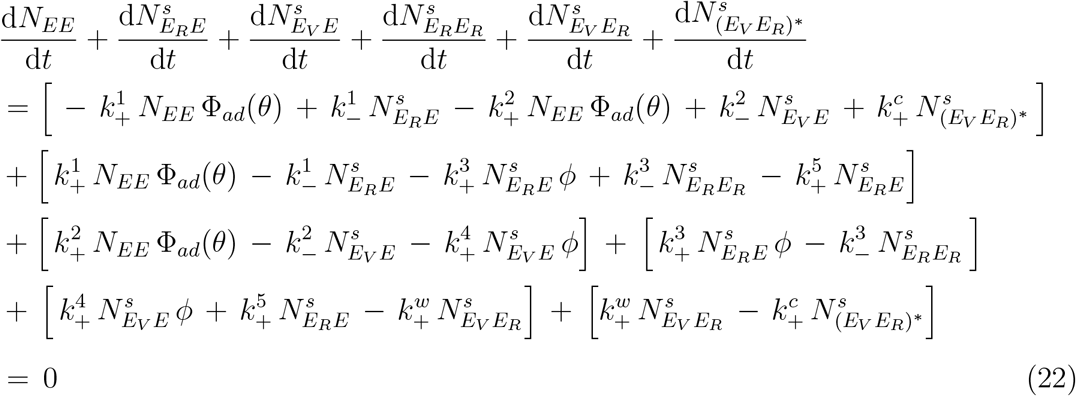

#### Balance for total number of lattice sites

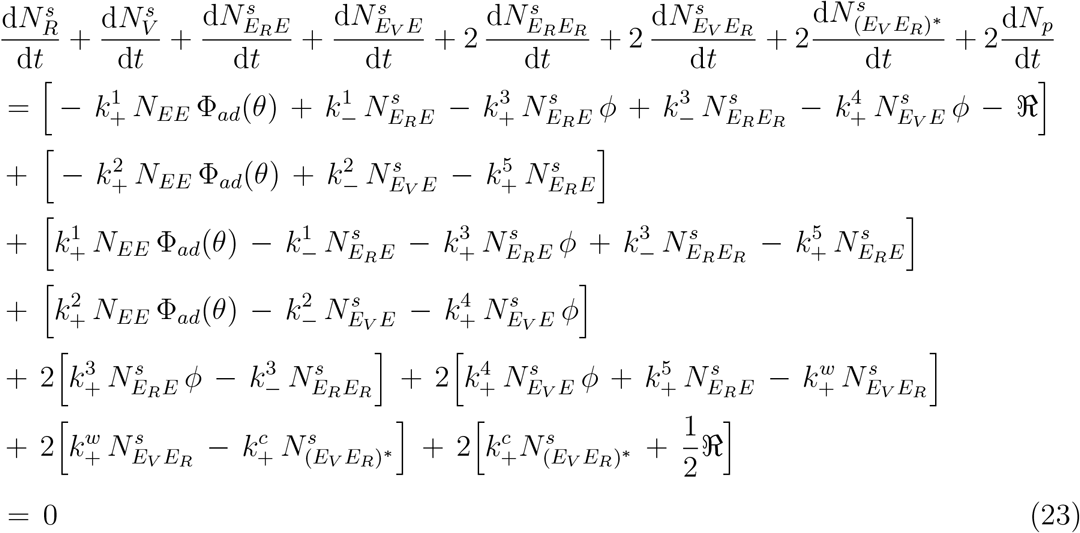

### SI5 Force-assisted removal and description related to ℜ

We assumed that the energy barrier for temporary winding-unwinding of a lattice site due to thermal fluctuations is symmetric in absence of enzymes. Both the rates to cross the energy barrier from either side are proportional to exp (−*E*_*m*_*/*(*k*_*B*_*T*)) resulting in zero net rate, where *E*_*m*_ is the energy required to cross the barrier, *k*_*B*_ is the Boltzmann constant, and *T* is the temperature. Due to a force *F* emerging from enzyme-induced unwinding (Eckhard et al., 2011), the energy barrier for other regular sites can become asymmetric. The energy required for the transition towards unwinding is reduced to (*E*_*m*_ − *λ*_*m*_*F*), and for the other side, it is increased to (*E*_*m*_ + *λ*_*m*_*F*). Here *λ*_*m*_ is the extent of dissociation in unwound state which is chosen as ∼ 3.6 Å (Perumal et al., 2008). Thus the net rate of flow over the energy barrier towards unwinding according to the theory of reaction rates is (Glasstone et al., 1941; Stuart and Anderson, 1953)

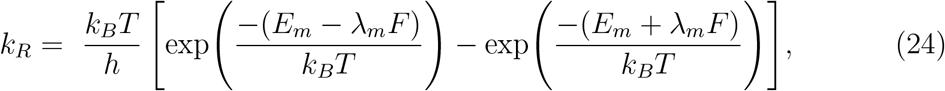

where *h* is Planck constant. For *F* = 0, the form (24) yields to zero. Here the value of *E*_*m*_ is chosen as *E*_*m*_ = *n*_*res*_ Δ*G/N*_*A*_, where Δ*G* = 1.9 kJ/mole per amino acid residue, *N*_*A*_ is Avogadro number, and *n*_*res*_ = 28 is the number of amino acid residues corresponding to one lattice site (Perumal et al., 2008). Thus the rate of force assisted removal of exposed regular sites is

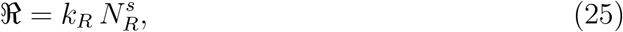

which is added in an *ad hoc* manner in the system of ODEs (Eqn (13)-(21)). We derive the force *F* in a heuristic manner.

The force corresponding to the local stress generated during enzyme-induced unwinding at one lattice (vulnerable) site is equivalent to the force necessary to generate a new surface via slippage of the chain on the surface (Adjari et al., 1994; Sung, 1995; Vega et al., 2001), i.e.

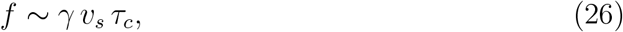

where 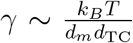 is the surface energy per unit area (Raphael and De Gennes, 1992; Léger and Creton, 2008), 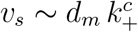 is the approximate velocity of slippage, and *τ*_*c*_ is a characteristic time. This characteristic time can be equivalent to the characteristic time of reptation of a polymer chain. The velocity of slippage *v*_*s*_ is approximately the multiplication of the length of the lattice site *d*_*m*_ to the frequency a chain experiences slippage events after cleavage i.e. 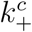. Note that the form for *f* in Eqn (26) is due to one unwound vulnerable site in presence of one enzyme. Due to multiple unwound vulnerable sites (in presence of enzymes), the total average force can be proportional to the rate at which enzymatic unwinding happens, i.e., 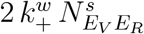(see Eqn (6)). Thus the average force per remaining lattice sites can be

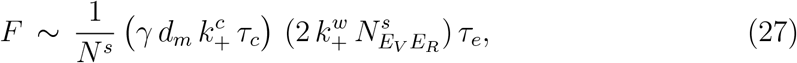

where *τ*_*e*_ is an average time required to form disentangled chains during detachment from the surface. When a fibril loses its lattice sites in degradation, dangling and entangled chains of the cleaved tropocollagen units (weakly attached to the fibril surface) appear continuously. In phenomenological manner and following the work of De Gennes (1979), we propose 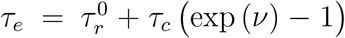 (Adjari et al., 1994; Vega et al., 2001), where *ν* is the number of lattice sites lost in form of disentangled chains during degradation. When *ν* is zero, i.e. in absence of degradation, 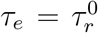 can be treated as average relaxation time of a fibril. The value of 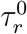 is 7 s (Shen et al., 2011; Gautieri et al., 2012). We treated *τ*_*c*_ as a material parameter and obtained its value as 1.4 s using the experimental data of Flynn et al. (2013) single fibril degradation under zero loading condition (see section SI10 and Fig. S2a).

### SI6 Reaction rate constants and other parameters

We provided the values of rate constants and other parameters in table S1. Using the relation of protein adsorption-desorption rate constants (Adamczyk, 2000), we set 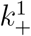 and 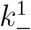 where 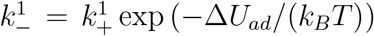. We assumed that all unbinding events happen with same probability (backwards reactions of Eqn (3) and (4)), and the corresponding rate constants are equal, i.e. 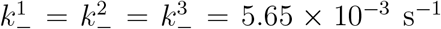 (Ottl et al., 2000). Using the value of 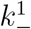 and for a chosen potential barrier difference of 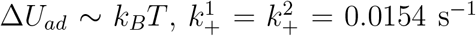. In experiments, the velocity of the enzyme on the collagen surface reported was ∼ 4.5 *×* 10^−6^ m/s (Saffarian et al., 2004). Using this reported value of the velocity, we set the forward rate constants of Eqn (4) and (5) related to full-length binding/hopping. The minimum distance an enzyme can traverse to find another lattice site for full-length binding or jump to change track is *d*_TC_ = 1.5 nm, which sets 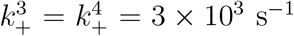. However, as the probability of the occurrence represented by the second part of Eqn (5) (see main text) is expected to be lower compared to the other hopping events, we set the value of 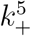 in an approximate manner based on the minimum distance between two neighboring vulnerable sites which is *D*_*gap*_ = 67 nm, and it sets 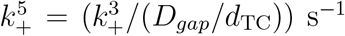. Using the transition state theory, we set the rate constant for enzyme-induced irreversible unwinding Eqn (6) as 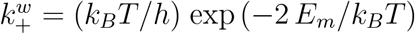, where *E*_*m*_ = *n*_*res*_ Δ*G/N*_*A*_, and (*E*_*V*_ *E*_*R*_)^*∗*^ corresponds to two lattice sites. For Δ*G* = 1.9 kJ/mole per amino acid residue (Perumal et al., 2008), *n*_*res*_ = 28 for one lattice site, *N*_*A*_ = 6.023 *×* 10^23^ and a chosen temperature *T* = 310 K, it yields to 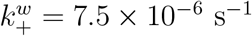. The value of 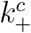 is different for different types of collagenase and collagen substrate (Welgus et al., 1982; Fields et al., 1987; Mallya et al., 1992; Lauer-Fields et al., 2000; Salsas-Escat et al., 2010). To test the present model, we used two values: 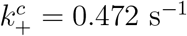 (fibroblast collagenase and native collagen type I) (Welgus et al., 1982) and 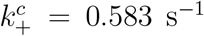(class I clostridium histolyticum collagenases and rat type I) (Mallya et al., 1992).

**Table S1:** List of parameters and their values.

| Parameter | Value |
| --- | --- |
| $d_{TC}$ | 1.5 nm (Chung et al., 2004) |
| $\ell_{TC}$ | 300 nm (Chung et al., 2004) |
| D-band gap ($D_{gap}$) | 67 nm (Chung et al., 2004) |
| $d_E$ | 10 nm (Tyn and Gusek, 1990) |
| $d_m$ | 8 nm (Perumal et al., 2008) |
| $n_{res}$ | 28 (Perumal et al., 2008) |
| $c_1$ | 25 (discussed in main text) |
| $\Delta G$ | 1.9 kJ/mole<br>per amino acid residue<br>(Perumal et al., 2008) |
| $\lambda_m$ | 3.6 Å(Perumal et al., 2008) |
| $k_+^1 = k_+^2$ | 0.0154 s<sup>-1</sup> (Adamczyk, 2000) |
| $k_-^1 = k_-^2 = k_-^3$ | 5.65 × 10<sup>-3</sup> s<sup>-1</sup> (Ottl et al., 2000) |
| $k_+^3 = k_+^4$ | 3 × 10<sup>3</sup> s<sup>-1</sup> (Saffarian et al., 2004) |
| $k_+^5$ | ($k_+^3/(D_{gap}/d_{TC})$) |
| $k_+^w$ | 7.5 × 10<sup>-6</sup> s<sup>-1</sup> Perumal et al. (2008) |
| $k_+^c$ | 0.583 s<sup>-1</sup> (Welgus et al., 1982)<br>or, 0.472 s<sup>-1</sup> (Mallya et al., 1992) |
| $\tau_r^0$ | 7 s (Shen et al., 2011) |
| $\tau_c$ | 1.4 s (obtained using experiment<br>of Flynn et al. (2013)) |

### SI7 Initial conditions to solve the ODEs

We used MATLAB ODE23 solver to solve the ODEs using the following initial conditions. The nine first order ODEs (Eqn (13)-(21)) require nine following initial conditions: at time *t* = 0, number of enzymes surrounding a fibril 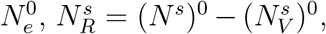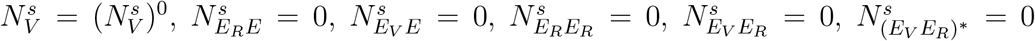 and *N*_*P*_ = 0. For a fibril of initial diameter 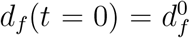 and length 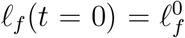, we obtained (*N* ^*s*^)^0^ and 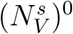 using Eqn (2) and Eqn (1), respectively.

### SI8 Fibril fraction estimation in simulation box from collagen concentration

To set the fibril fraction *ϕ*_*f*_ (the ratio of the volume of all collagen fibrils to the volume of the simulation box) in our simulations, we followed a crude analytical way converting the collagen concentration to *ϕ*_*f*_. An example is the following. We consider a collagen concentration ∼ 2.5 mg/mL (Ranamukhaarachchi et al., 2019). Note that we performed *in vitro* experiments using the same collagen concentration. For this chosen concentration and using the molecular weight of tropocollagen ∼ 300 kDa (León-López et al., 2019), the number of tropocollagen units in the simulation box volume (which is 125 µm^3^) is ∼ 0.63 *×* 10^6^. The volume of one tropocollagen is 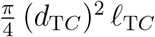. Thus the total volume of all tropocollagen molecules in the simulation box volume is 334 *×* 10 µm^3^.

These tropocollagen molecules arrange and generate fibrils in a solvent medium. Because of the gaps in between tropocollagen molecules in a fibril in hydrated condition (which is difficult to estimate in the present work), the total volume of all fibrils must be higher than the total volume of all tropocollagen molecules. To obtain a nearly correct estimate of the total volume of all fibrils, we assumed packing factor for cylindrical tropocollagen in loosely packed condition as 0.4 (note that the value of packing factor for spheres in loosely packed state is in the range 0.4 − 0.56). Thus the nearly corrected volume of all fibrils is (334 *×* 10^−3^ µm^3^)*/*0.4, and this volume yields to a fibril fraction *ϕ*_*f*_ ∼ 0.006 for the chosen collagen concentration 2.5 mg/mL.

The collagen percentage in different organs (such as, brain, liver, heart, kidney, lung and colon, etc.) varies in the range 0.1 − 6% (Tarnutzer et al., 2023). We performed simulations with different values of *ϕ*_*f*_ in the range 0.003 − 0.03, which is 0.3 − 3%.

## SI9 Experimental methods

### Chemicals

High concentration, rat tail acid extracted type I collagen was procured from Corning (Corning, NY). PEG (8000 Da) was ordered in powder form from Sigma-Aldrich (St Louis, MO) and reconstituted in PBS (Life Technologies, Carlsbad, CA) immediately before usage with a final concentration of 100 mg/mL, and 1× reconstitution buffer was composed of sodium bicarbonate, HEPES free acid, and nanopure water.

### Preparation of collagen gels with different microstructures

First, PEG of required amounts to make a 2 or 8 mg/mL final concentration (denoted as P2 or P8) was added to the DMEM. This is followed by addition of the reconstitution buffer and mixing. Thereafter, the collagen stock was added to the mixture to produce a final concentration of 2.5 mg/mL. Finally, pH of the final mixture was adjusted using 1 N NaOH, followed by incubation (∼ 45 minutes) at 37°C. Following polymerization, PEG was washed out of the gels by rinsing with the DMEM (3× for 5 minutes each). For collagenase treatment, gels were incubated with bacterial collagenase of 2.5 µg/mL for about 45 minutes. (We added 50 microliters of 10 microgram per mL of collagenase on top of the gels which yields final concentration of collagen to be 2.5 µg/mL.)

### Fast green staining and imaging of collagen gels

The prepared collagen gels were fixed with 4% paraformaldehyde for 30 mins. After that, the gels were washed thoroughly at least 3 times by subjecting them to shaking and replacing with PBS for 10 min. The gels were then incubated with 100 µg/mL of fast green dye in PBS (Fast green FCF, Thermo Fischer, USA) and were subjected to shaking overnight. The gels were washed with PBS at least 3 times in an orbital shaker for a duration of 30 minutes each time. The stained gels were then imaged using a confocal fluorescence microscope (Leica, SP8) with 40x water immersion objective. Fast green was excited at a wavelength of 627 nm, and 630-730 nm was used for detection.

### Scanning electron microscopy (SEM)

SEM was performed on the gels using FEI SEM Apreo equipped with ETD detector. Both the collagenase treated gels and the non-treated gels were first fixed with 4% paraformaldehyde for 1 hour. This is followed by 3x rinsing in PBS for 10 minutes in each step of shaking. The gels were then rinsed twice with Milli-Q water for 15 minutes each. The fixed gels were then subjected to a series of dehydration steps in ethanol and hexamethyldisilazane (HMDS) using the protocol outlined in Raub et al. (2007). Briefly, the gels were dehydrated first in ethanol dilution series: 30%, 50%, 70%, 90% and 100% for 15 minutes of each step. The gels were then incubated in ethanol/HMDS dilution series: 33%, 50%, 66% and 100% for 15 minutes each. After the final incubation, the gels were allowed to dry on an aluminum foil for at least 1 day in the fume hood. The dried gel samples were then sputter coated with a Pelco SC-7 sputter coater with gold as the target. The gels were then imaged at 5 kV and 0.6 nA with magnifications of 15000x and 30000x.

## SI10 Single fibril model validation

We used the experimental data of Flynn et al. (2013) on single fibril degradation to calibrate and validate our single fibril model. Flynn et al. (2013) reported the diameter of degrading single collagen fibril under different external loading condition in a 5 µM Clostridium histolyticum bacterial collagenase type A solution. To solve our single fibril model for the parameter sets reported in Flynn et al. (2013), we need to obtain the number of enzymes 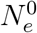 for a fibril. Using their enzyme concentration, we estimated 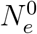 for a fibril in an approximate manner based on a volume element equivalent to the initial volume of the fibril. For fibrils of initial diameter 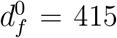 and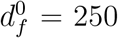, the values of 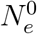 are 2100 and 750, respectively, for a chosen length 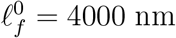.

The value of *τ*_*c*_ = 1.4 s is obtained by calibrating our model prediction to the single fibril data corresponding to 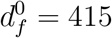 under zero load condition (see Fig. 3a). The value of 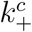 used to calibrate our model is 0.583 s^−1^ (class I clostridium histolyticum collagenases and rat type I) (Mallya et al., 1992). In a nutshell, our single fibril model have satisfactorily captured the experimental findings of Flynn et al. (2013) (see Fig. 3a).

### The choice of *δE*_*m*_ = 0.013 *E*_*m*_

The degradation rate reduces when the fibril is under external tension (Bhole et al., 2009; Camp et al., 2011; Flynn et al., 2013) (see Fig. 3a). Perhaps the external tension increases the stability of the triple helices in the fibril by increasing the energy barrier for enzymatic unwinding (Chang et al., 2012; Chang and Buehler, 2014; Tonge et al., 2015; Saini et al., 2020; Topol et al., 2021). However, the cleavage mechanism can be different for the isolated tropocollagen (triple helix) molecules under external tension (Adhikari et al., 2011, 2012). For a fibril, the increase in the internal energy due to external loading can increase the energy barrier for enzymatic unwinding where the rate is proportional to 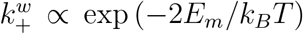. The increase in the energy barrier results in the decrease in the rate of unwinding, implying a decrease in the rate of degradation.

It is difficult to provide a direct estimation of the increase in the energy barrier of enzymatic unwinding because of external loading. We have followed a simple analytical approach to obtain the increment in the energy following the work of Tonge et al. (2015) and using a general expression for strain energy (purely elastic). Under external loading, we can write the increment in energy Δ*U* for a tropocollagen under low external load *f*_*ext*_ = 2 pN is

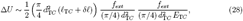

where *δ𝓁* is the extension and *E*_T*C*_ is the elastic modulus of a tropocollagen. The value of *E*_T*C*_ is in the range 0.01 − 1 GPa (David et al., 1978; Sun et al., 2002; Gautieri et al., 2012). For a chosen value *E*_T*C*_ ∼ 0.5 GPa and *δ𝓁* in the range 50 − 250 nm (Sun et al., 2002), (28) yields to Δ*U* ∼ 0.009 *E*_*m*_ − 0.015 *E*_*m*_, where *E*_*m*_ = *n*_*res*_ Δ*G/N*_*A*_. Our model captured the experimental trend (Flynn et al., 2013) well for *δE*_*m*_ = 0.013 *E*_*m*_.

### The scaling related to single fibril degradation

Figs. 3b,c imply that the degradability must be directly proportional to the number of enzymes per unit surface area of the fibril. According to the model assumptions, if the enzymes are not loosing their activity and potency as enzymatic degradation progresses, then the surface area of the fibril is the only parameter which decreases with time. Thus a thicker fibril and a thinner fibril of same length can degrade up to a same extent if the ratio 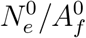, where 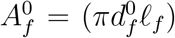 is the area of the fibril, is fixed. This is reflected in Fig. S2a-c, where the same extent of degradation is observed if 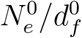 is a constant. The model predicts thicker fibrils with significantly larger number of enzymes to be highly degradable than thinner fibrils with very less number of enzymes subject to the 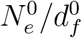, and *viceversa*. The predictions are sensitive to the choice of 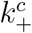 which varies with different MMPs and types of collagen. As expected, the extent of degradation decreases with the decrease in 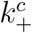.

## SI11 Model microstructure generation for P2 and P8 gels using the histograms of fibril diameters from the experiments

The lognormal distribution curves fit our histrograms better than other distributions such as normal and gamma distributions (see Fig. S4).

We used the lognormal fitted curves of the diameter histograms (P2 and P8) from the experiments to generate the fibril distributions. The parameters related to the lognormal fits from P2 and P8 are *µ*_*P* 2_ = −2.247 and *σ*_*P* 2_ = 0.45, and *µ*_*P* 8_ = −2.288 and *σ*_*P* 8_ = 0.4, respectively. Using these values and lognormal function, random numbers are generated in between 0.04-0.2 µm for P2, 0.03-0.16 µm for P8, and we ignore the tail regions where PDF is less than 2 (see Fig. S5a). In both P2 and P8, all fibrils are of same length. For P2, *𝓁*_*f*_ */l*_*p*_ = 2, and *𝓁*_*f*_ */l*_*p*_ = 1.8 for P8 which is 10% smaller than that of P2. We note in passing that in experiments, the mean length of P8 is found to be slightly (approx. 10 %) smaller than that of P2. For both P2 and P8, the distributions are generated such that the volume fraction of fibrils *ϕ*_*f*_ = 0.007 turns out to be the same. The outcomes of number of fibrils *n*_*f*_ in P2 and P8 for simulation are 40 and 75, respectively. The chosen value of *ϕ*_*f*_ is close to the collagen concentration used in the experiments 2.5 mg/mL. For each P2 and P8, we performed three independent simulations, and we generated all three sets using the histograms of diameters from experiments (Fig. S5a). We showed the thickness distributions of the generated fibrils in Fig. S5b. The distributions generated for the simulations are qualitatively similar to those from the experiments.

**Figure S2:**
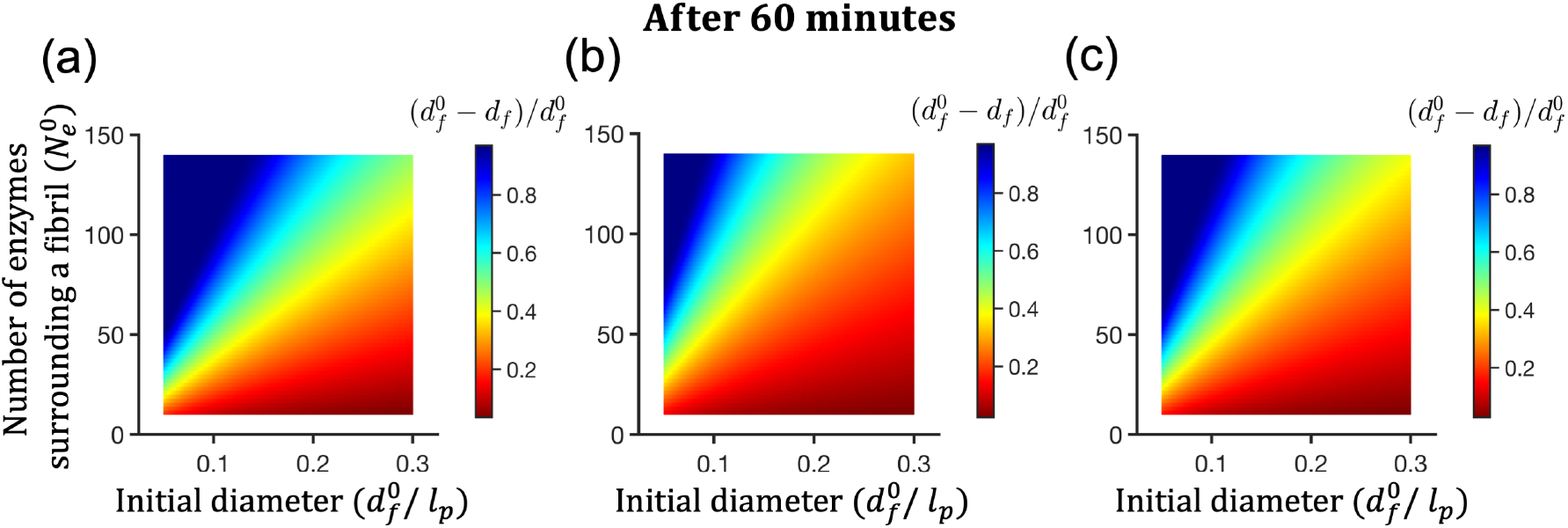
Degradation of single collagen fibril. (a)-(c) are color maps representing the extent of degradation after 1 hour for ranges of 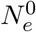 (number of enzymes surrounding a fibril) and 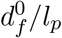 (initial fibril diameter) where *l*_*p*_ = 1 *µ*m is a nominal length scale. (a) fibril length 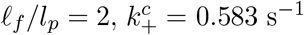 (Mallya et al., 1992); (b) fibril length 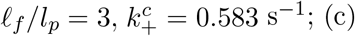 fibril length 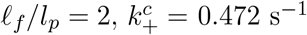 (Welgus et al., 1982).

**Figure S3:**
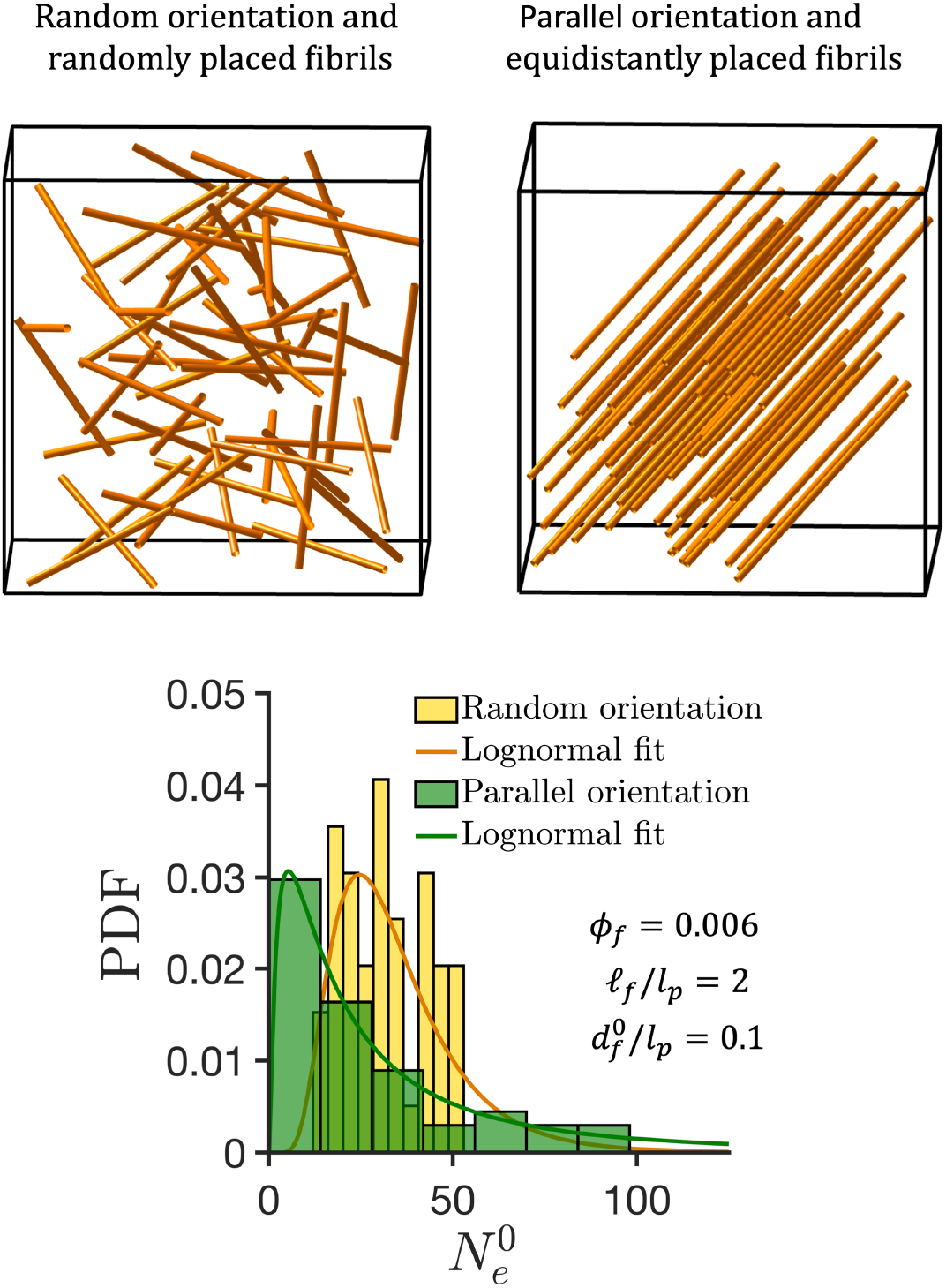
The enzyme distributions in two different configurations.

**Figure S4:**
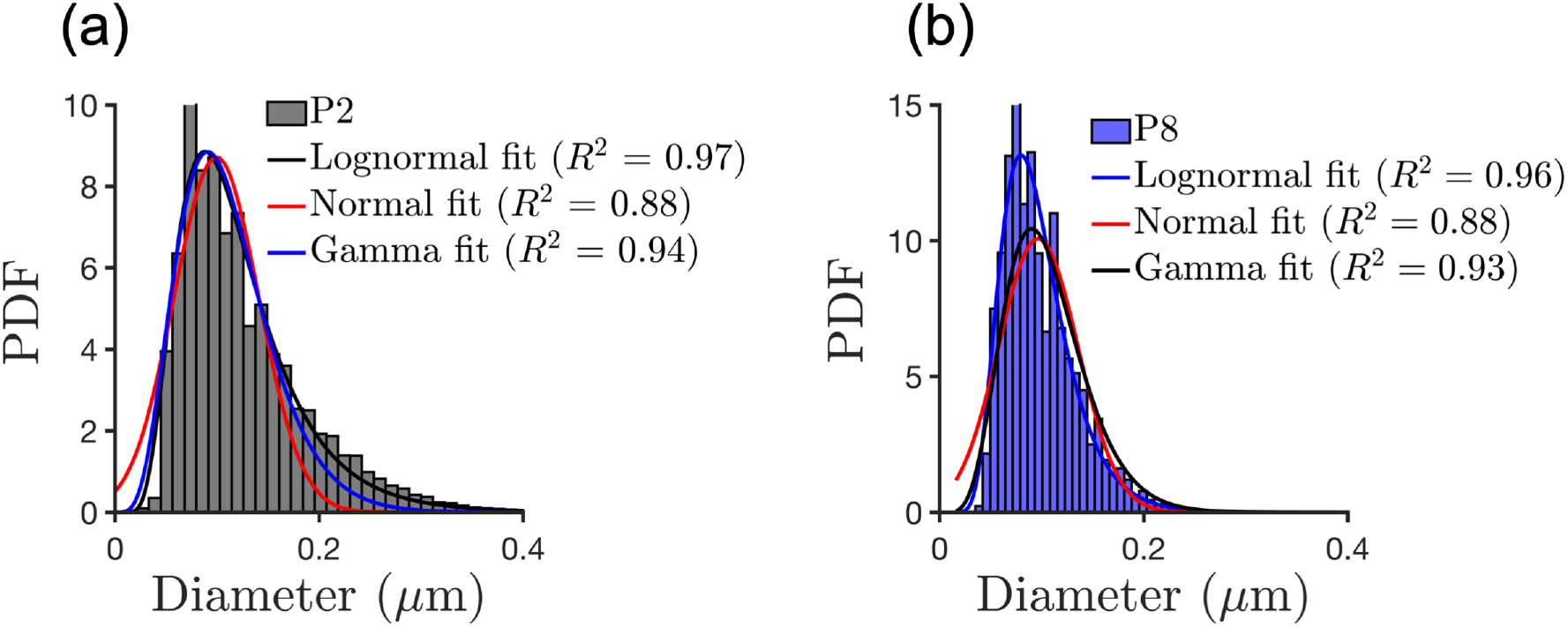
Different fits and goodness of fits for P2 (a) and P8 (b)

**Figure S5:**
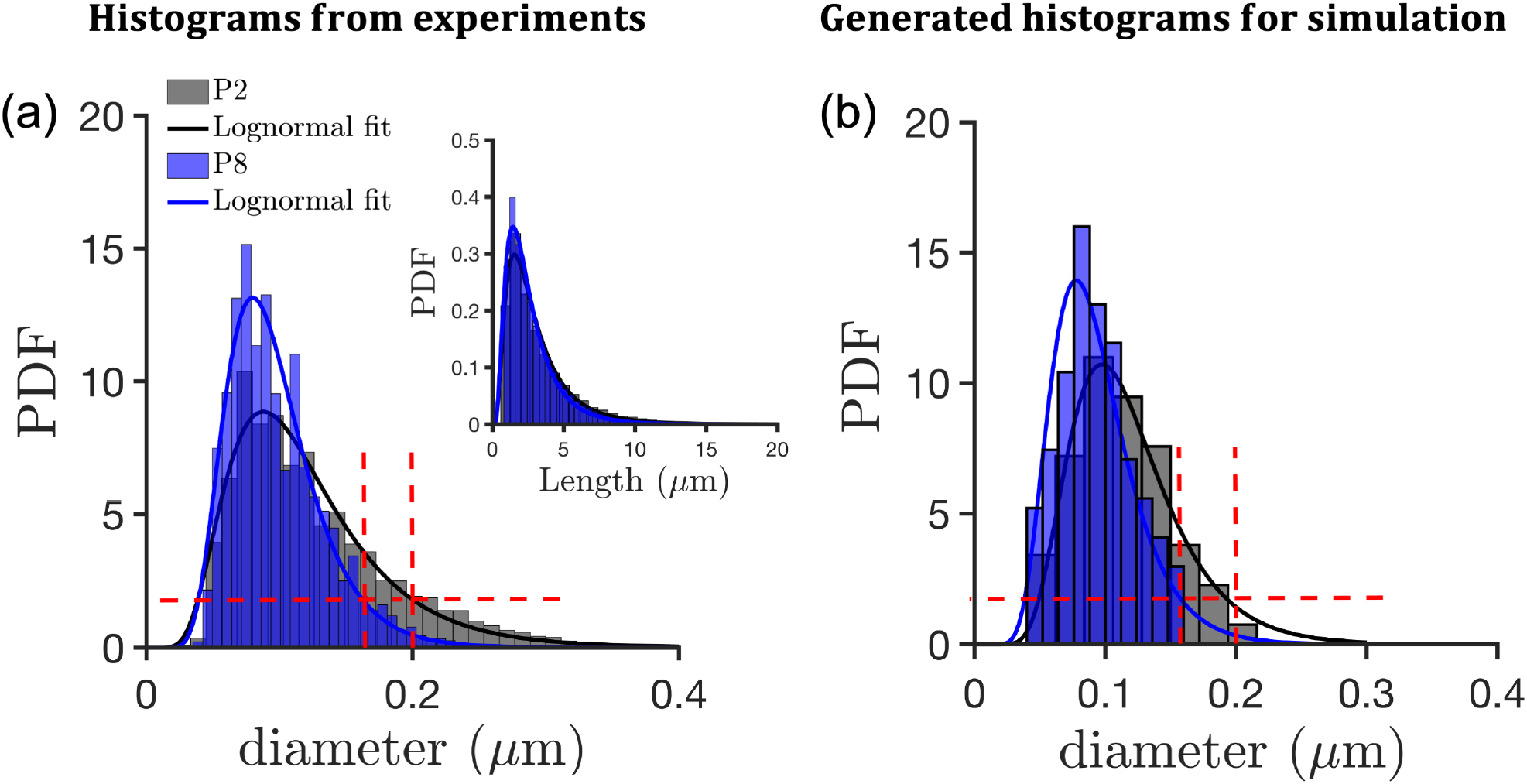
Histograms of diameters for P2 and P8 matrices. (a) From experiments. (b) Histograms are generated for hybrid simulation using (a)

## References

Adamczyk, Z. (2000). Kinetics of diffusion-controlled adsorption of colloid particles and proteins. JCIS, 229(2).

Adamczyk, Z., Senger, B., Voegel, J., and Schaaf, P. (1999). Irreversible adsorption/deposition kinetics: A generalized approach. J. Chem. Phys., 110(6).

Adhikari, A., Chai, J., and Dunn, A. (2011). Mechanical load induces a 100-fold increase in the rate of collagen proteolysis by mmp-1. J. American Chem. Soc., 100:513a.

Adhikari, A., Glassey, E., and Dunn, A. (2012). Conformational dynamics accompanying the proteolytic degradation of trimeric collagen i by collagenases. J. American Chem. Soc., 134:13259–13265.

Adjari, A., Brochard-Wyart, F., de Gennes, P., Leibler, L., Viovy, J., and Rubinstein, M. (1994). Slippage of an entangled polymer melt on a grafted surface. Physica A: Statistical Mechanics and its Applications, 204:17–39.

Ashworth, J. and Cox, T. (2024). The importance of 3d fibre architecture in cancer and implications for biomaterial model design. Nat. Rev. Cancer, 24:461–479.

Ayad, A. (2024). Generate fiber. https://www.mathworks.com/matlabcentral/fileexchange/73392-generate-fiber.

Bhole, A., Flynn, B., Liles, M., Saeidi, N., Dimarzio, C., and Ruberti, J. (2009). Mechanical strain enhances survivability of collagen micronetworks in the presence of collagenase: implications for load-bearing matrix growth and stability. Phil. Tran. Roy. Soc. A: Mathematical, Physical and Engineering Sciences, 367:3339–3362.

Buehler, M. (2008). Nanomechanics of collagen fibrils under varying cross-link densities: atomistic and continuum studies. JMBBM, 1(1).

Camp, R., Liles, M., Beale, J., Saeidi, N., Flynn, B., Moore, E., Murthy, S., and Ruberti, J. (2011). Molecular mechanochemistry: low force switch slows enzymatic cleavage of human type i collagen monomer. J. American Chem. Soc., 133(11).

Chang, S. and Buehler, M. (2014). Molecular biomechanics of collagen molecules. Materials Today, 17:70–76.

Chang, S., Flynn, B., Ruberti, J., and Buehler, M. (2012). Molecular mechanism of force induced stabilization of collagen against enzymatic breakdown. Biomaterials, 33:3852–3859.

Chen, P., Chen, X., Hepfer, R., Damon, B., Shi, C., Yao, J., Coombs, M., Kern, M., Ye, T., and Yao, H. (2021). A noninvasive fluorescence imaging-based platform measures 3d anisotropic extracellular diffusion. Nat. Commun., 12:1913.

Chung, L., Dinakarpandian, D., Yoshida, N., Lauer-Fields, J., Fields, G., Visse, R., and Nagase, H. (2004). Collagenase unwinds triple-helical collagen prior to peptide bond hydrolysis. EMBO, 23:3020–3030.

Cichocki, B. and Hinsen, K. (1990). Dynamic computer simulation of concentrated hard sphere suspensions: I. simulation technique and mean square displacement data. Physica A, 166(3).

Daly, A., Riley, L., Segura, T., and Burdick, J. (2020). Hydrogel microparticles for biomedical applications. Nat. Rev. Mat., 5:20–43.

David, L., Grood, E., Noyes, F., Zernicke, R., et al. (1978). Biomechanics of ligaments and tendons. Exercise and sport sciences reviews, 6:125–182.

De Gennes, P. (1979). Scaling concepts in polymer physics. Cornell university press.

Dewavrin, J., Hamzavi, N., Shim, V., and Raghunath, M. (2014). Tuning the architecture of three-dimensional collagen hydrogels by physiological macromolecular crowding. Acta Biomat., 10:4351–4359.

Eckhard, U., Schönauer, E., Nüss, D., and Brandstetter, H. (2011). Structure of collagenase g reveals a chew-and-digest mechanism of bacterial collagenolysis. Nat. Struct. Mol. Biol., 18:1109–1114.

Ermak, D. and McCammon, J. (1978). Brownian dynamics with hydrodynamic interactions. J. Chem. Phys., 69(4).

Eyring, H. (1935). The activated complex and the absolute rate of chemical reactions. Chem. Rev., 17(1).

Eyring, H. (1936). Viscosity, plasticity, and diffusion as examples of absolute reaction rates. J. Chem. Phys., 4(4).

Fang, F., Satulovsky, J., and Szleifer, I. (2005). Kinetics of protein adsorption and desorption on surfaces with grafted polymers. Biophys. J., 89(3).

Fields, G., Van Wart, H., and Birkedal-Hansen, H. (1987). Sequence specificity of human skin fibroblast collagenase. evidence for the role of collagen structure in determining the collagenase cleavage site. J. Biol. Chem., 262(13).

Flynn, B., Tilburey, G., and Ruberti, J. (2013). Highly sensitive single-fibril erosion assay demonstrates mechanochemical switch in native collagen fibrils. BMMB, 12:291–300.

Gautieri, A., Vesentini, S., Redaelli, A., and Buehler, M. (2012). Viscoelastic properties of model segments of collagen molecules. Matrix Biology, 31:141–149.

Glasstone, S., Laidler, K., and Eyring, H. (1941). The theory of rate processes: the kinetics of chemical reactions, viscosity, diffusion and electrochemical phenomena. McGrawHill Book, New York.

Gupta, V., Vaishnavi, V., Arrieta-Ortiz, M., Ps, A., Km, J., Jeyasankar, S., Raghunathan, V., Baliga, N., and Agarwal, R. (2024). 3d hydrogel culture system recapitulates key tuberculosis phenotypes and demonstrates pyrazinamide efficacy. Adv. Healthcare Mater., page 2304299.

Han, S., Makareeva, E., Kuznetsova, N., DeRidder, A., Sutter, M., Losert, W., Phillips, C., Visse, R., Nagase, H., and Leikin, S. (2010). Molecular mechanism of type i collagen homotrimer resistance to mammalian collagenases. J. Biol. Chem., 285(29).

Huang, L., Haylor, J., Hau, Z., Jones, R., Vickers, M., Wagner, B., Griffin, M., Saint, R., Coutts, I., El Nahas, A., et al. (2009). Transglutaminase inhibition ameliorates experimental diabetic nephropathy. Kidney International, 76:383–394.

Islam, M., Tudryn, G., and Picu, C. (2016). Microstructure modeling of random composites with cylindrical inclusions having high volume fraction and broad aspect ratio distribution. Compt. Mat. Sci., 125:309–318.

Iyer, S., Visse, R., Nagase, H., and Acharya, K. (2006). Crystal structure of an active form of human mmp-1. J. Mol. Biol., 362(1).

Kim, S., Min, S., Choi, Y., Jo, S., Jung, J., Han, K., Kim, J., An, S., Ji, Y., Kim, Y., et al. (2022). Tissue extracellular matrix hydrogels as alternatives to matrigel for culturing gastrointestinal organoids. Nat. Commun., 13:1692.

Lauer-Fields, J., Tuzinski, K., Shimokawa, K., Nagase, H., and Fields, G. (2000). Hydrolysis of triple-helical collagen peptide models by matrix metalloproteinases. J. Biol. Chem., 275(18).

Léger, L. and Creton, C. (2008). Adhesion mechanisms at soft polymer interfaces. Phil. Tran. Roy. Soc. A, 366(1869).

León-López, A., Morales-Peñaloza, A., Martínez-Juárez, V., Vargas-Torres, A., Zeugolis, D., and Aguirre-Álvarez, G. (2019). Hydrolyzed collagen—sources and applications. Molecules, 24:4031.

Liotta, L., Tryggvason, K., Garbisa, S., Hart, I., Foltz, C., and Shafie, S. (1980). Metastatic potential correlates with enzymatic degradation of basement membrane collagen. Nature, 284:67–68.

Mallya, S., Mookhtiar, K., and Van Wart, H. (1992). Kinetics of hydrolysis of type i, ii, and iii collagens by the class i and ii clostridium histolyticum collagenases. J. Protein Chem., 11.

Manka, S., Carafoli, F., Visse, R., Bihan, D., Raynal, N., Farndale, R., Murphy, G., Enghild, J., Hohenester, E., and Nagase, H. (2012). Structural insights into triple-helical collagen cleavage by matrix metalloproteinase 1. Proc. Nat. Acad. Sci., 109(31).

Marino, M. and Vairo, G. (2014). Stress and strain localization in stretched collagenous tissues via a multiscale modelling approach. Computer Methods in Biomechanics and Biomedical Engineering, 17:11–30.

McKleroy, W., Lee, T., and Atabai, K. (2013). Always cleave up your mess: targeting collagen degradation to treat tissue fibrosis. American J. Physiology-Lung Cellular and Molecular Physiology, 304:L709–L721.

Metzmacher, I., Radu, F., Bause, M., Knabner, P., and Friess, W. (2007). A model describing the effect of enzymatic degradation on drug release from collagen minirods. Euro. J. Pharma. Biopharma., 67(2):349–360.

Nagase, H. and Visse, R. (2011). Triple helicase activity and the structural basis of collagenolysis. In Extracellular Matrix Degradation, pages 95–122. Springer.

Narasimhan, B. and Fraley, S. (2024). Degradability tunes ecm stress relaxation and cellular mechanics. BioRxiv, pages 2024–07.

Nebuloni, M., Albarello, L., Andolfo, A., Magagnotti, C., et al. (2016). Insight on colorectal carcinoma infiltration by studying perilesional extracellular matrix. Sci. Rep., 6:22522.

Nerenberg, P., Salsas-Escat, R., and Stultz, C. (2008). Do collagenases unwind triplehelical collagen before peptide bond hydrolysis? reinterpreting experimental observations with mathematical models. Proteins, 70(4).

Okada, T., Hayashi, T., and Ikada, Y. (1992). Degradation of collagen suture in vitro and in vivo. Biomaterials, 13(7).

Orgel, J., Irving, T., Miller, A., and Wess, T. (2006). Microfibrillar structure of type i collagen in situ. Proc. Nat. Acad. Sci., 103(24).

Ottl, J., Gabriel, D., Murphy, G., Knäuper, V., Tominaga, Y., Nagase, H., Kröger, M., Tschesche, H., Bode, W., and Moroder, L. (2000). Recognition and catabolism of synthetic heterotrimeric collagen peptides by matrix metalloproteinases. Chemistry & Biology, 7(2).

Perumal, S., Antipova, O., and Orgel, J. (2008). Collagen fibril architecture, domain organization, and triple-helical conformation govern its proteolysis. Proc. Nat. Acad. Sci., 105(8).

Philp, C., Siebeke, I., Clements, D., Miller, S., Habgood, A., John, A., Navaratnam, V., Hubbard, R., Jenkins, G., and Johnson, S. (2018). Extracellular matrix cross-linking enhances fibroblast growth and protects against matrix proteolysis in lung fibrosis. American J. Respiratory cell and Molecular Biology, 58:594–603.

Ranamukhaarachchi, S., Modi, R., Han, A., Velez, D., Kumar, A., Engler, A., and Fraley, S. (2019). Macromolecular crowding tunes 3d collagen architecture and cell morphogenesis. Biomaterials Sci., 7(2).

Raphael, E. and De Gennes, P. (1992). Rubber-rubber adhesion with connector molecules. J. Phys. Chem., 96(10).

Raub, C., Suresh, V., Krasieva, T., Lyubovitsky, J., Mih, J., Putnam, A., Tromberg, B., and George, S. (2007). Noninvasive assessment of collagen gel microstructure and mechanics using multiphoton microscopy. Biophys. J., 92:2212–2222.

Ray, N., van Noorden, T., Radu, F., Friess, W., and Knabner, P. (2013). Drug release from collagen matrices including an evolving microstructure. ZAMM-Journal of Applied Mathematics and Mechanics, 93(10-11).

Saffarian, S., Collier, I., Marmer, B., Elson, E., and Goldberg, G. (2004). Interstitial collagenase is a brownian ratchet driven by proteolysis of collagen. Science, 306(5693).

Saini, K., Cho, S., Dooling, L., and Discher, D. (2020). Tension in fibrils suppresses their enzymatic degradation–a molecular mechanism for ‘use it or lose it’. Matrix Biol., 85.

Salsas-Escat, R., Nerenberg, P., and Stultz, C. (2010). Cleavage site specificity and conformational selection in type i collagen degradation. Biochemistry, 49(19).

Sarkar, S., Marmer, B., Goldberg, G., and Neuman, K. (2012). Single-molecule tracking of collagenase on native type i collagen fibrils reveals degradation mechanism. Current Biol., 22(12).

Schaaf, P. and Talbot, J. (1989). Surface exclusion effects in adsorption processes. J. Chem. Phys., 91(7).

Schultz, K. and Anseth, K. (2013). Monitoring degradation of matrix metalloproteinases-cleavable peg hydrogels via multiple particle tracking microrheology. Soft Matter, 9(5).

Seo, B., Chen, X., Ling, L., Song, Y., Shimpi, A., Choi, S., Gonzalez, J., et al. (2020). Collagen microarchitecture mechanically controls myofibroblast differentiation. Proc. Nat. Acad. Sci., 117:11387–11398.

Shen, Z., Kahn, H., Ballarini, R., and Eppell, S. (2011). Viscoelastic properties of isolated collagen fibrils. Biophys. J, 100:3008–3015.

Smith, S., Cianci, C., and Grima, R. (2017). Macromolecular crowding directs the motion of small molecules inside cells. J. Roy. Soc. Interface, 14(131).

Smith, S. and Grima, R. (2017). Fast simulation of brownian dynamics in a crowded environment. J. Chem. Phys., 146(2).

Staunton, J., Vieira, W., Fung, K., Lake, R., Devine, A., and Tanner, K. (2016). Mechanical properties of the tumor stromal microenvironment probed in vitro and ex vivo by in situ-calibrated optical trap-based active microrheology. Cellular and Molecular Bioengineering, 9:398–417.

Stuart, D. and Anderson, O. (1953). Dependence of ultimate strength of glass under constant load on temperature, ambient atmosphere, and time. J. American Ceramic Soc., 36(12).

Stylianopoulos, T., Diop-Frimpong, B., Munn, L., and Jain, R. (2010). Diffusion anisotropy in collagen gels and tumors: the effect of fiber network orientation. Biophys. J., 99:3119–3128.

Sun, Y., Luo, Z., Fertala, A., and An, K. (2002). Direct quantification of the flexibility of type i collagen monomer. Biochem. Biophys. Res. Commun., 295:382–386.

Sung, W. (1995). Slippage of linear flows of entangled polymers on surfaces. Phys. Rev. E, 51:5862.

Talbot, J., Tarjus, G., Van Tassel, P., and Viot, P. (2000). From car parking to protein adsorption: an overview of sequential adsorption processes. Colloids and Surfaces A, 165(1-3).

Tarnutzer, K., Siva Sankar, D., Dengjel, J., and Ewald, C. (2023). Collagen constitutes about 12% in females and 17% in males of the total protein in mice. Sci. Rep., 13:4490.

Tobolsky, A. and Eyring, H. (1943). Mechanical properties of polymeric materials. J. Chem. Phys., 11(3).

Tonge, T., Ruberti, J., and Nguyen, T. (2015). Micromechanical modeling study of mechanical inhibition of enzymatic degradation of collagen tissues. Biophysical J., 109:2689–2700.

Topol, H., Demirkoparan, H., and Pence, T. (2021). Fibrillar collagen: A review of the mechanical modeling of strain-mediated enzymatic turnover. Applied Mech. Rev., 73:050802.

Tyn, M. and Gusek, T. (1990). Prediction of diffusion coefficients of proteins. Biotech. and Bioengg., 35(4).

Tzafriri, A., Bercovier, M., and Parnas, H. (2002). Reaction diffusion model of the enzymatic erosion of insoluble fibrillar matrices. Biophysical journal, 83(2).

Vega, D., Villar, M., Alessandrini, J., and Valles, E. (2001). Terminal relaxation of model poly (dimethylsiloxane) networks with pendant chains. Macromolecules, 34:4591–4596.

Visse, R. and Nagase, H. (2003). Matrix metalloproteinases and tissue inhibitors of metalloproteinases: structure, function, and biochemistry. Circulation Res., 92(8).

Vuong, A., Rauch, A., and Wall, W. (2017). A biochemo-mechano coupled, computational model combining membrane transport and pericellular proteolysis in tissue mechanics. Proc. Royal Soc. A, 473(2199).

Welgus, H., Jeffrey, J., Stricklin, G., and Eisen, A. (1982). The gelatinolytic activity of human skin fibroblast collagenase. J. Biol. Chem., 257(19).

Winkler, J., Abisoye-Ogunniyan, A., Metcalf, K., and Werb, Z. (2020). Concepts of extracellular matrix remodelling in tumour progression and metastasis. Nat. Commun., 11:5120.

Wohlgemuth, R., Brashear, S., and Smith, L. (2023). Alignment, cross linking, and beyond: a collagen architect’s guide to the skeletal muscle extracellular matrix. American J. Physiology-Cell Physiology, 325:C1017–C1030.

Wynn, T. and Ramalingam, T. (2012). Mechanisms of fibrosis: therapeutic translation for fibrotic disease. Nat. Medicine, 18:1028–1040.

